# Perivascular SPP1 Mediates Microglial Engulfment of Synapses in Alzheimer’s Disease Models

**DOI:** 10.1101/2022.04.04.486547

**Authors:** Sebastiaan De Schepper, Judy Z Ge, Gerard Crowley, Laís SS Ferreira, Dylan Garceau, Christina E Toomey, Dimitra Sokolova, Thomas Childs, Tammaryn Lashley, Jemima J Burden, Steffen Jung, Michael Sasner, Carlo Sala Frigerio, Soyon Hong

**Author notes:** Corresponding Author: Dr. Soyon Hong, Gower St, London WC1E 6BT, United Kingdom.

## Abstract

Microglia are phagocytes of the brain parenchyma, where they interact with neurons to engulf synapses in a context-dependent manner. Genetic studies in Alzheimer’s disease (AD) highlight dysfunctional phagocytic signaling in myeloid cells as disease-associated pathway. In AD models, there is a region-specific reactivation of microglia-synapse phagocytosis involving complement; however, what drives microglia-synapse engulfment remains unknown. Here, we show that SPP1 (Osteopontin), a glycoprotein associated with inflammation, is regionally upregulated and modulates microglial synaptic engulfment in AD mouse models. Ultrastructural examination revealed SPP1 expression predominantly by perivascular macrophages, a subtype of border-associated macrophages, in the hippocampus of mice and patient tissues. Cell-cell interaction networks of single-cell transcriptomics data suggested that perivascular SPP1 drives microglial functional states in the hippocampal microenvironment of AD mice. Absence of *Spp1* expression resulted in failure of microglia to mediate synaptic phagocytosis. This study suggests a critical role for perivascular SPP1 in neuroimmune crosstalk in AD-relevant context.

## Introduction

Neuronal and synaptic homeostasis critically relies on the integrated contribution of neuroimmune signaling, which is governed by resident immune cells that continuously monitor and respond to signals in the surrounding brain parenchyma^1–3^. Microglia are the primary resident macrophages of the central nervous system (CNS) that contribute to activity-dependent circuit refinement, in part by mediating engulfment of synaptic elements^4^. In contrast to microglia, the role of perivascular macrophages (PVM), a subtype of border-associated macrophages that reside in the perivascular space of the CNS juxtaposed with the brain parenchyma, is less well characterized^5^. PVM share their primitive yolk-sac origin with microglia and occupy the vascular niche as early as E10.5, where they likely are the primary responders to toxic agents and pathogens and regulate blood-brain barrier (BBB) permeability and transport of nutrients and metabolites^6–8^. Their strategic location adjacent to blood vessels may allow PVM to act as a regulated gateway to the CNS parenchyma^9^. Microglia and PVM reside in functionally distinct but bordering compartments, raising the question of whether these two CNS-resident macrophages communicate with each other to coordinate neuronal and synaptic health in physiological and disease states.

Genetic studies in late-onset Alzheimer’s disease (AD), which reflect about 90 % of AD cases, pinpoint at dysfunctional phagocytosis as key disease-associated processes^10,11^. More than half of the risk factors identified in AD are expressed by CNS-resident macrophages, including PVM and microglia, and converge on endolysosomal and phagocytic pathways. Microglia in AD models have been shown to engulf synapses, thereby potentially contributing to synaptic pathology that represents a significant correlate of cognitive impairment in AD^12,13^. A key mechanism by which microglia mediate phagocytosis of synapses is via a region-specific activation of the classical complement cascade involving C1q^4,12,14^. Genetic and antibody-mediated inhibition of the classical complement cascade leads to reduction of microglia-synapse engulfment, and ultimately, amelioration of synaptic loss and cognitive decline in multiple AD models^12,15–17^. However, what reactivates the region-specific engulfment of synapses by microglia in the AD hippocampus is unknown. Given the genetic rationale and accumulative functional data on neuroimmune signaling as key modulators in AD, mechanistic and functional insight into phagocytic-centered dysfunction of microglia and environmental factors that imprint their fate will be critical to identify therapeutic and biomarker targets to assess and alter AD pathogenesis.

An interesting candidate to this end is secreted phosphoprotein 1 (SPP1/ Osteopontin). In multiple organs throughout the body, SPP1 has been shown to be expressed by macrophage subsets, T cells and other immune cells to coordinate various macrophage functions, including phagocytosis^18–21^. In the brain throughout lifespan, *Spp1* expression is spatiotemporally upregulated in macrophages residing in specific regions that have a high and local requirement for tissue remodeling and phagocytosis, e.g., axon tract-associated microglia in postnatal corpus callosum, tissue-repairing microglia in ischemic regions after stroke, and activated-response and disease-associated microglia/macrophages surrounding amyloid-β (Aβ) plaques in aged AD brains^22–26^. Collectively, this suggests that SPP1 in myeloid cells in the brain is associated with transient phagocytic requirements in line with what has been suggested in peripheral tissues^20^. Interestingly, levels of secreted SPP1 are increased in the cerebrospinal fluid (CSF) and plasma of early AD, correlating with cognitive decline^27–30^. Further, SPP1^+^ macrophage subsets have been suggested to be engaged in synapse phagocytosis in the developing brain^23^. This altogether raises the question of whether SPP1 is involved in microglia-synapse phagocytosis in AD, and if so, how. Yet, the cellular origin and the functional relevance of SPP1 in AD pathogenesis are unknown.

Here, we report that SPP1 is required for synaptic phagocytosis by microglia. Notably, super- and ultrastructural examination in the hippocampus revealed that *Spp1* is expressed predominantly by PVM, and to a lesser extent, perivascular fibroblasts (PVF). Furthermore, we show that SPP1 is upregulated early in a region-specific manner in multiple AD mouse models. Specifically, in hAPP knock-in (*App*^NL-F^) mice, SPP1 induction occurs at a timepoint prior to robust plaque deposition, but when hippocampal synapses are already vulnerable to microglial engulfment and oligomeric Aβ (oAβ) is found accumulating on the vasculature. Genetic ablation of SPP1 ameliorates oAβ-induced microglial C1q activation as well as synaptic engulfment by microglia in multiple mouse models. NicheNet post single-cell transcriptomics revealed multiple autocrine and paracrine signaling pathways modulated by SPP1, with many converging on potential microglial functional states. Altogether, our data suggest that microglial phagocytic states are triggered by perivascular SPP1 signaling in pre-plaque hippocampus of *App*^NL-F^ mice when synapses are vulnerable. We propose a functional role for perivascular SPP1 in modulating microglial phagocytosis of synapses, thereby impeding microglia-synapse interactions in AD.

## Results

### Perivascular SPP1 upregulation coincides with complement activation and microglia-synapse engulfment in pre-plaque *App*^NL-F^ hippocampus

To address whether, and if so, where and when, SPP1 production is dysregulated early in AD^27–30^, we utilized the slow-progressing *App*^NL-F^ mouse model, where control by the endogenous *App* promoter allows for physiological cell-type specific and temporal regulation of Aβ production^31^. We first assessed whether microglia engulf synapses in a region-specific manner in the *App*^NL-F^ hippocampus, akin to what has been described in other AD mouse models^12,32^. We analyzed synaptic Homer1^+^ immunoreactive synaptic puncta within CD68^+^ microglial lysosomes in *App*^NL-F^ hippocampus at the age of 6-month-old (6 mo) when plaques have not developed yet (Fig. 1A, Suppl Fig. 1A). We observed an approximate sevenfold increase in Homer1^+^ synapses engulfed and localized within lysosomes of P2Y12^+^ microglia in *App*^NL-F^ mice or wildtype (WT) mice, suggesting increased levels of microglial phagocytosis (Fig. 1B). Coinciding with synaptic phagocytosis in 6 mo *App*^NL-F^ brains, we observed a region-specific increase of C1q protein deposition, the initiating protein of the classic complement cascade (Fig. 1C,D). Of note, *C1qa* expression in the *App*^NL-F^ mice was contained within *Tmem119*^+^ microglia as assessed by single-molecule fluorescent *in-situ* hybridization (smFISH), suggesting that microglia are producers of C1q in the 6 mo *App*^NL-F^ hippocampus, in line with earlier reports (Fig. 1C, insert)^12,33^. As previously reported^31^, we did not see overt Aβ plaque deposition in the brain parenchyma of 6 mo *App*^NL-F^ hippocampus, in contrast to 15 mo *App*^NL-F^ brains that showed overt plaque deposition (Suppl Fig. 1A). These data suggest that there is a region-specific upregulation of microglial C1q and synaptic engulfment in pre-plaque 6 mo *App*^NL-F^ hippocampus akin to earlier studies in pre-plaque brains of more aggressive transgenic mouse models of AD^12^. We next assessed expression levels of SPP1 in the 6 mo *App*^NL-F^ hippocampus. Stimulated emission depletion (3D-*τ*-STED) super-resolution imaging revealed approximately threefold increase of punctate SPP1 protein immunoreactivity in the CA1 hippocampus of *App*^NL-F^ mice as compared to age- and sex-matched WT controls (Fig. 1E,F). The SPP1 upregulation was region-specific, i.e., in hippocampus but not cerebellum (Suppl Fig. 1B, C), as measured by immunohistochemistry (IHC). In line with increased SPP1 production, we found an approximate threefold increase of *Spp1* mRNA expression levels in hippocampal CA1 sections of *App*^NL-F^ mice compared to WT mice by smFISH in intact tissue, further confirmed by qPCR analysis on brain homogenates (Fig. 1G,H). The specificity of SPP1 antibody and *Spp1-*targeting smFISH probes was validated by absence of signals in *Spp1*^*KO/KO*^ mice (Suppl Fig. 1D). These data altogether establish that SPP1 is upregulated in the hippocampus of 6 mo *App*^NL-F^ mice at the onset of microglial C1q deposition and synaptic engulfment.

**Figure 1.**
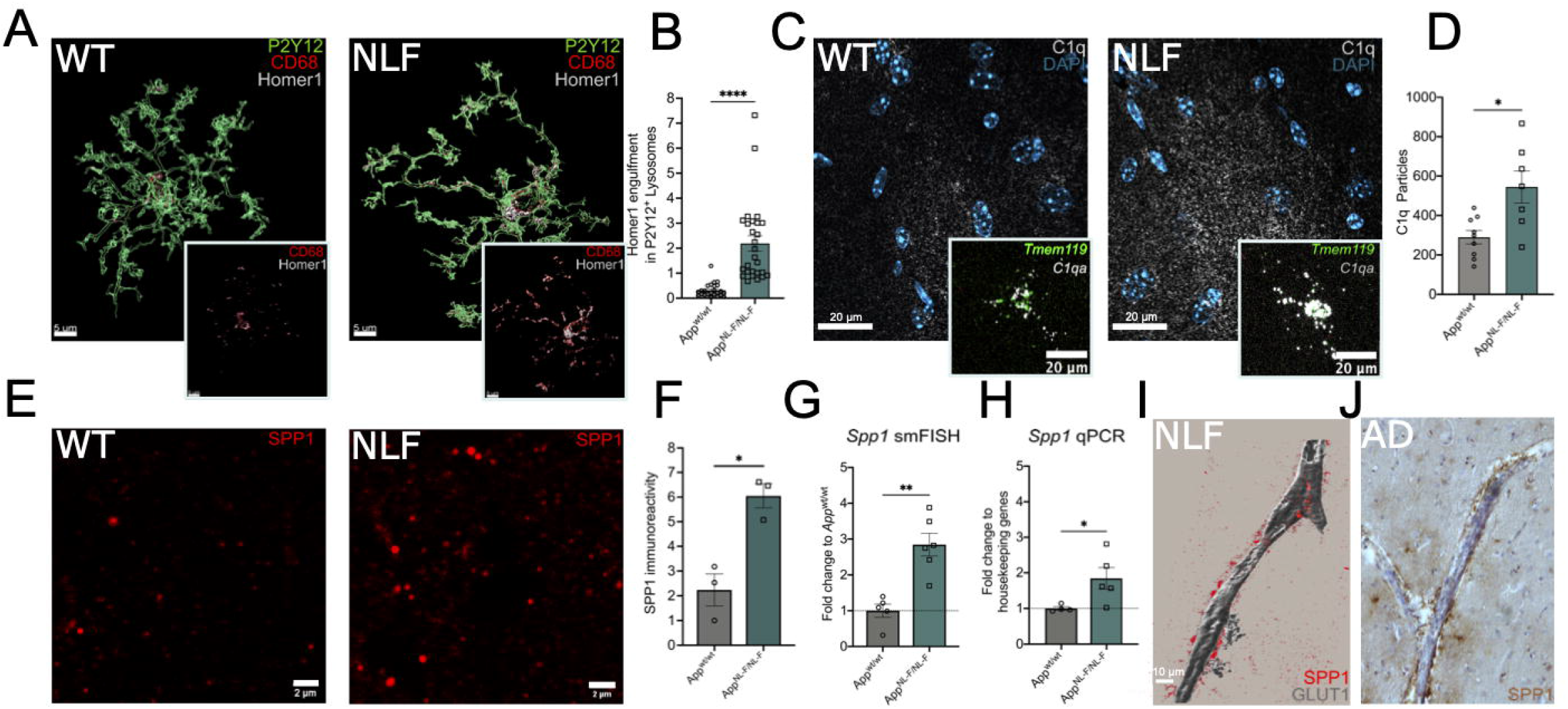
Perivascular SPP1 upregulation coincides with complement activation and microglia-synapse engulfment in pre-plaque *App*^NL-F^ hippocampus. **(A)** Representative 3D reconstructed images showing Homer1 engulfment within CD68^+^ lysosomes of P2Y12^+^ microglia in 6 mo *App*^WT^ (WT) versus *App*^NL-F^ CA1 hippocampal *Stratum Lacunosum Moleculare* (SLM). Scale bar represents 5 µm. **(B)** Quantification of Homer1 engulfment in 6 mo WT and *App*^NL-F^ P2Y12^+^ microglia. Two-tailed unpaired Student’s t-test, n=13-25 cells total, n=4 mice pooled from 2 independent experiments. **(C)** Representative confocal images of C1q protein expression in 6 mo WT versus *App*^NL-F^ mice. Insert represents *C1qa* mRNA within *Tmem119*^*+*^ microglia as identified by single-molecule fluorescent *in-situ* hybridization (smFISH). Data are representative of at least 5 independent experiments. Scale bar represents 20 µm. **(D)** Quantification of C1q particles (puncta) in 6 mo WT and *App*^NL-F^ CA1 hippocampus. Mann-Whitney test, n=7-9 mice from 2 independent experiments. **(E**,**F)** Representative stimulated emission depletion (3D-*τ*-STED) imaging of secreted SPP1 in 6 mo WT and *App*^NL-F^ SLM **(E)** and quantification **(F)**. Mann-Whitney test, n=3. Scale bar represents 2 µm. **(G**,**H)** Quantification of *Spp1* expression within hippocampus of 6 mo WT versus *App*^NL-F^ as measured by smFISH on intact hippocampal tissue **(G)** or via qPCR on whole hippocampal homogenates **(H)**. Mann-Whitney test, n=3, 2 independent experiments. **(I)** Representative 3D reconstruction of SPP1 adjacent to GLUT1^+^ blood vessels in SLM of 6 mo *App*^NL-F^. Scale bar represents 10 µm. **(J)** Representative image of SPP1 expression along vasculature in post-mortem hippocampal brain slices of AD patients. See also Suppl Table 1. *p < 0.05 **p < 0.01 and ****p < 0.0001. Data are shown as Mean ± SEM.

Notably, we found that *Spp1* mRNA expression was particularly enriched in cells within the *Stratum Lacunosum Moleculare* (SLM) layer of the hippocampal CA1 and that *Spp1*^*+*^ cells displayed an elongated vascular-like pattern (Suppl Fig. 1E). Further co-staining with pan-endothelial marker GLUT1 indeed showed cellular zonation of *Spp1* mRNA expression adjacent to vascular structures, whereas outside of the endothelial membrane, no *Spp1* mRNA was detected (Suppl Fig. 1E., insert) To confirm the vascular distribution of SPP1 on the protein level, we used confocal imaging and 3D surface rendering and confirmed close association of cytosolic SPP1 expression with GLUT1^+^ vasculature (Fig. 1I). Further, using volumetric confocal imaging analysis, we observed a notable deposition of oAβ along the vasculature of the 6 mo *App*^NL-F^ hippocampus (Suppl Fig. 1F), as assessed by the oAβ-specific NAB61^+^ immunostaining juxtaposed to GLUT1^+^ vasculature in hippocampal CA1 of 6 mo *App*^NL-F^ mice^34^. NAB61^+^ staining was not observed in hippocampus of 6 mo WT mice (Suppl Fig. 1F); we also observed a similar increase of Aβ immunoreactivity using 4G8 and HJ3.4 antibodies, which recognize Aβ17-24 and Aβ1-13, respectively, along GLUT1^+^ vasculature (Suppl Fig. 1G). Further, oAβ deposition along vasculature was more pronounced in 15 mo *App*^NL-F^ mice, in contrast to age-matched WT controls in which no positive NAB61 staining was found (Suppl Fig. 1H). Importantly, also high-resolution confocal imaging in hippocampal brain tissue of AD patients showed a striking presence of SPP1 immunoreactivity along the vasculature (Fig. 1J, Suppl Table 1), similar to what we observed in the murine *App*^NL-F^ hippocampus. Altogether, our data show that SPP1 is increased along the vasculature in *App*^NL-F^ mice and in human AD, and that this upregulation occurs prior to robust plaque deposition, when complement and synapse pathology in the hippocampus become evident in *App*^NL-F^ mice.

### SPP1 marks perivascular macrophages and fibroblasts in steady-state and pre-plaque *App*^NL-F^ hippocampus when synapses are being engulfed

We next sought to determine the cellular source of SPP1 in 6 mo *App*^NL-F^ hippocampus, where *Spp1* upregulation was restricted to the perivascular space (Fig. 1). Given the role of SPP1 as a marker of macrophage subsets in peripheral tissues, we first assessed whether *Spp1* is expressed by CNS-resident macrophages, i.e., *Tmem119*^*hi*^ P2Y12^+^ microglia and *Cd163*^*+*^ CD206^+^ perivascular macrophages (PVM) that associate with the basement membrane. To this end, we performed smFISH in combination with IHC (smFISH-IHC) and high-resolution confocal imaging to ensure cell-type specific volumetric assessment of mRNA. Notably, we did not find any appreciable *Spp1* expression in *Tmem119*^hi^ P2Y12^+^ microglia. Rather, *Spp1* signal was predominantly restricted to *Tmem119*^lo/neg^P2Y12^-^ cells in the *App*^NL-F^ hippocampi (Suppl Fig. 2A), suggesting that microglia are not a major source of *Spp1* in the pre-plaque AD hippocampus. Specifically, *Spp1* expression was localized to cells labelled for CD163 and the pan-PVM marker CD206, that were closely associated with GLUT1^+^ vasculature (Fig. 2A,B). We further observed *Spp1* co-expression with PVM-specific platelet factor 4 (*Pf4*) and *Cd163* in the hippocampus of 6 mo *App*^NL-F^ mice (Suppl Fig. 2C;)^5,35^. Interestingly, we also found some *Spp1*^*+*^ *Cd163*^*-*^ *Pf4*^*-*^ cells in the hippocampal perivascular space (Suppl Fig. 2C). *Spp1* mRNA has been suggested by single cell RNA-sequencing (scRNA-seq) to be expressed by vascular leptomeningeal cells (VLMC), also known as perivascular fibroblasts (PVF), which are located within parenchymal arteries^35^ (Suppl Fig. 2D). Indeed, the second cluster of *Spp1*^*+*^ cells co-expressed the pan-PVF marker *Pdgfra* (encoding CD140a), suggesting PVF as a second cellular source of SPP1 surrounding GLUT1^+^ vasculature in the hippocampus (Suppl Fig. 2E, Fig. 2C). Of note, *Spp1* expression in microglia was detected at later stages, i.e., in 6E10^+^ plaque-rich 15 mo *App*^NL-F^ hippocampus (Suppl Fig. 2F), likely reflecting SPP1^+^ disease-associated microglia^25^. These results establish that in WT and pre-plaque AD brains, SPP1 is produced mainly by vasculature-associated PVM and PVF, but not microglia.

**Figure 2.**
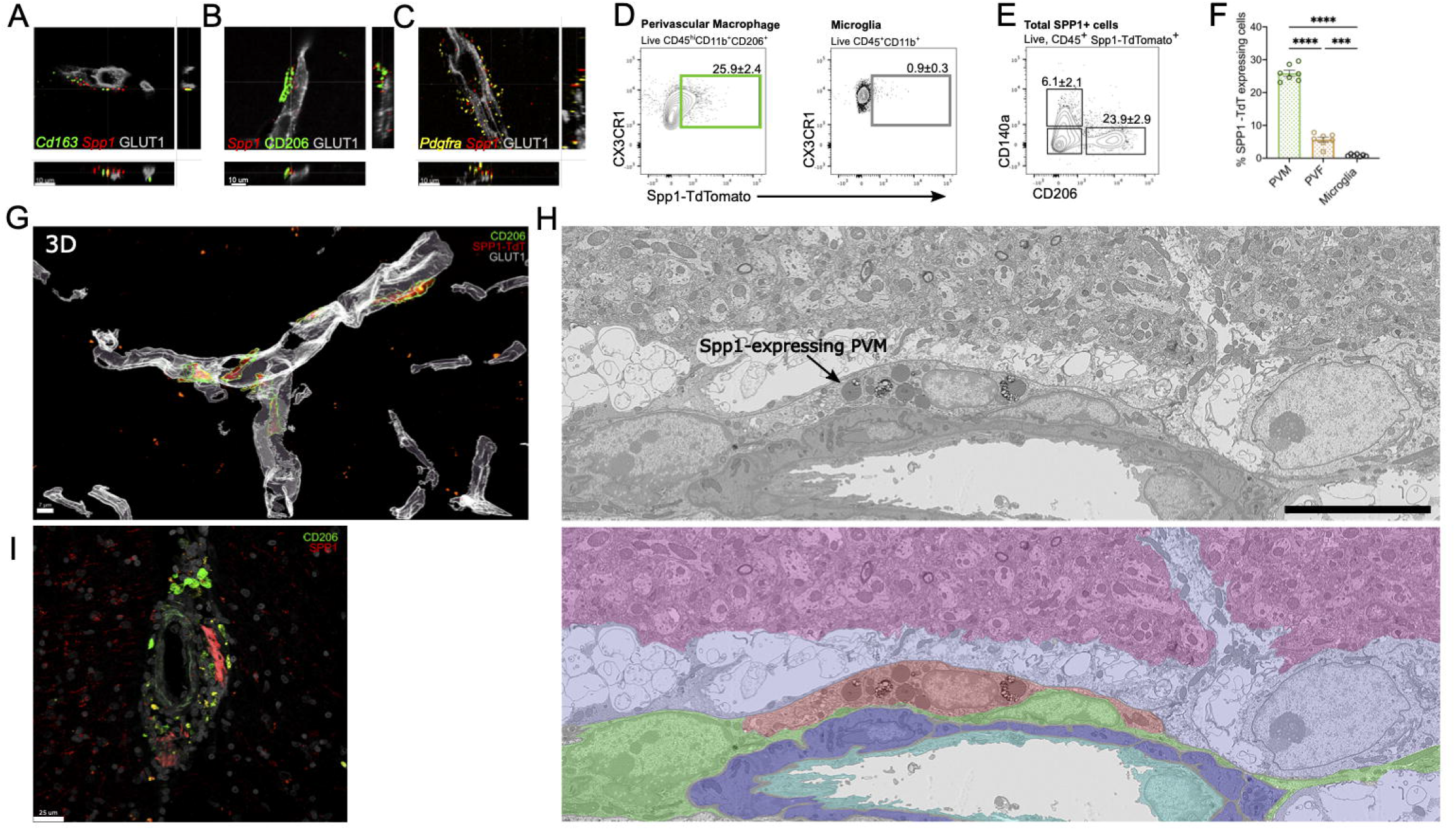
SPP1 marks perivascular macrophages and fibroblasts in steady-state and pre-plaque *App*^NL-F^ hippocampus when synapses are being engulfed. **(A-C)** Representative images of *Spp1* mRNA expression juxtaposed to GLUT1^+^ vasculature, colocalizing with pan-perivascular macrophage (PVM) markers *Cd163* **(A)**, CD206 **(B)** and perivascular fibroblasts (PVF, *Pdgfra*^+^) **(C)** in 6 mo *App*^NL-F^ SLM as characterized by smFISH-IHC. Scale bar represents 10µm. Data are representative of n=3 independent experiments. **(D-F)** Representative FACS plots to identify PVM (CXCR1^+^CD45^+^CD11b^+^CD206^+^, **D)**, microglia (CX3CR1^high^CD45^+^CD11b^+^, **E)** or gated on total SPP1-TdTomato (TdT) expressing cells isolated from *Spp1*^TdT^ hippocampal homogenates and quantification **(F)**. One way ANOVA, Kruskal-Wallis n=6 for all genotypes. Data are representative of n=2 independent experiments. **(G)** 3D reconstruction of CD206^+^ PVM expressing SPP1-Td along GLUT1^+^ vessels in SLM from *Spp1*^TdT^ mice. Scale bar represents 7 µm. Data are representative of n=2 independent experiments. **(H)** Representative single serial section SEM backscatter electron image of a representative SPP1-TdTomato positive PVM as identified by CLEM (Upper). SPP1-TdT-positive cell manually pseudo-coloured red, together with neuropil (pink), astrocytes (lilac), smooth muscle cells (purple), endothelial cells (cyan) and other perivascular cells (green), shown with reduced opacity over the electron microscopy data (Lower). Accompanying confocal overlays and correlation images shown in Suppl. 3E and 3D array tomography data shown in Suppl. Movie 1. Scale bar represents 10µm **(I)** Representative image of perivascular SPP1 in AD post-mortem hippocampal tissue, co-stained with CD206. Scale bar represents 25 µm. Data is representative of n=6, 6 different patient tissues (Suppl Table 1). ***p < 0.005 and ****p < 0.001. Data are shown as Mean ± SEM.

To further assess *in vivo* SPP1 expression, we developed *Spp1*-IRES-TdTomato (TdT) (*Spp1*^TdT^) reporter mice, which carry an IRES-TdT cassette in *Spp1* exon 7 retaining endogenous SPP1 expression (Suppl Fig. 3A-C). Flow cytometric analysis of naïve animals revealed the expression of *Spp1-*TdT within 25.9 % of pre-gated CD45^hi^ CD206^+^ PVM, in contrast to CD11b^+^ CD45^int^ CX3CR1^hi^ microglia that were almost devoid of TdT expression (0.9 % of which were TdT^+^) (Fig. 2D-F). Further, only approximately 6 % of total live TdT^+^ cells were CD140a^+^ (gene product of *Pdgfra*), in contrast to 23.9 % for CD206^+^, highlighting PVM as a cellular source of SPP1, although a third of TdT^+^ cells remains to-be-identified (Fig. 2E-F). Using high-resolution confocal imaging, we found that the distribution of SPP1-TdT was comparable to the SPP1-immunoreactivity detected by IHC (Fig. 1I), i.e., along GLUT1^+^ vasculature of the hippocampus (Suppl Fig. 3D). Further, we saw similar cellular localization of SPP1-TdT within CD206^+^ PVM in the *Spp1*^TdT^ hippocampus (Fig. 2G). Finally, we used correlative light and electron microscopy (CLEM) to target and visualize the ultrastructure and environmental context of SPP1-TdT positive cells in the hippocampus (Fig. 2H, Suppl Fig. 3E, Suppl Movie 1). CLEM identified SPP1-TdT expressing cells as lysosome-rich PVM located within the basement membrane of the perivascular space (Fig. 2H, Suppl Fig. 3E). CLEM and mRNA *in-situ* localization identified PVM as a source for SPP1 in mice, which translated to human tissue, where we found enrichment of SPP1 within the perivascular space of AD patients, occasionally overlaying with CD206^+^ cells (Fig. 2I). Collectively, our data suggest that CD206^+^*Cd163*^*+*^*Pf4*^*+*^ PVM and *Pdgfra*^*+*^/CD140^+^ PVF are the cellular source of Spp1 in hippocampus in mice and humans.

**Figure 3:**
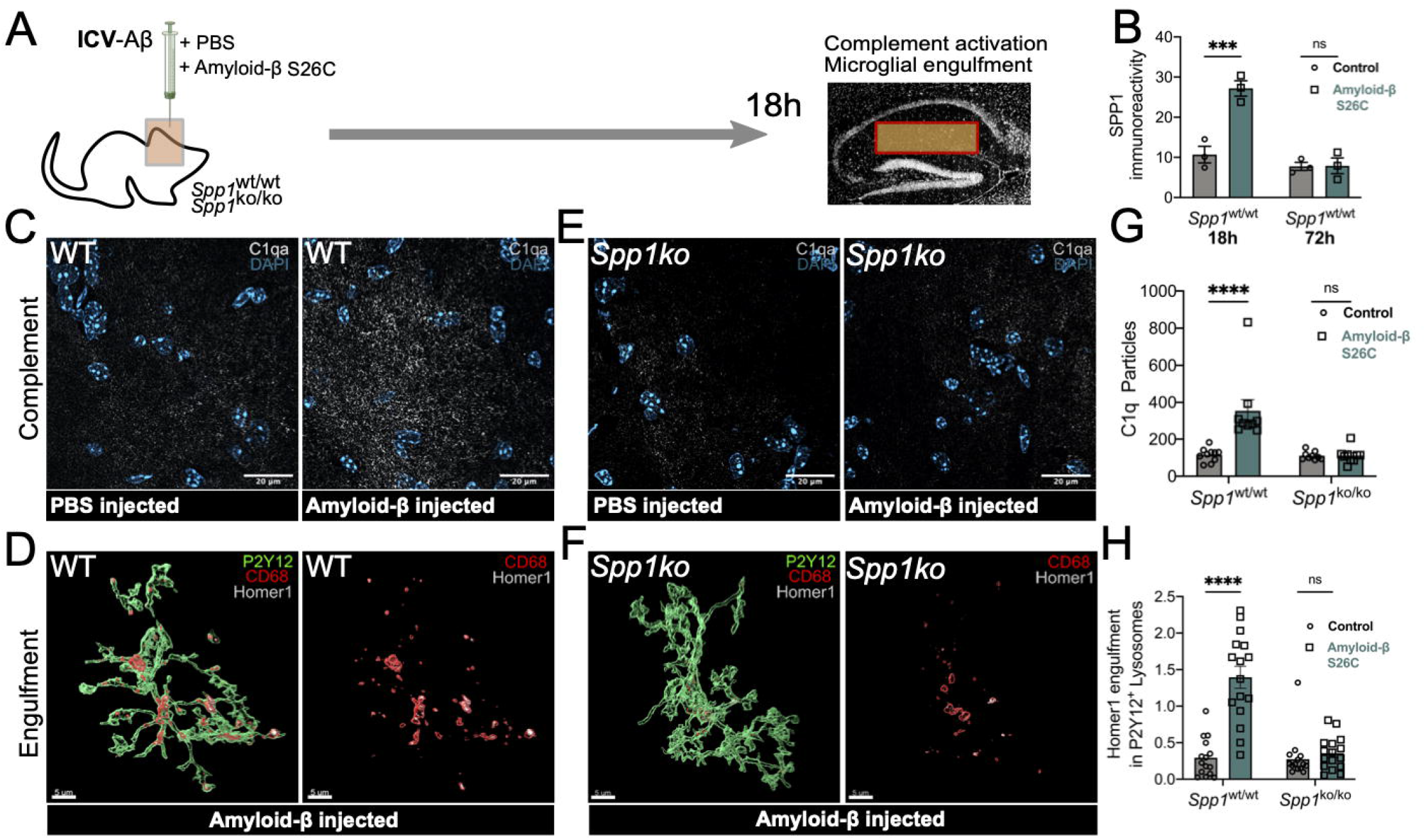
*Spp1* modulates complement activation and microglial synaptic engulfment upon acute oAβ challenge. **(A)** Scheme illustrating intracerebroventricular (ICV) injection of S26C oAβ versus PBS in WT versus *Spp1*^*KO/KO*^ mice, 18 h before tissue collection and analysis. **(B)** Quantification of SPP1 immunoreactivity within SLM of 3 mo WT mice injected with oAβ versus PBS control, at either 18 h or 72 h post-ICV injection. Two-way ANOVA, n=3 from 1 independent experiment. **(C)** Representative images of C1q expression in SLM of 3 mo PBS versus oAβ-injected WT mice **(D)** Representative 3D reconstructed images showing Homer1 engulfment within CD68^+^ lysosomes of P2Y12^+^ microglia from WT mice injected with oAβ. **(E)** Representative images of C1q expression in *Spp1*^*KO/KO*^ mice injected with PBS versus oAβ. **(F)** Representative 3D reconstructed images showing Homer1 engulfment within CD68^+^ lysosomes of microglia from *Spp1*^*KO/KO*^ mice injected with oAβ. **(G)** Quantification of C1q particles (puncta) in WT or *Spp1*^*KO/KO*^ mice treated with either PBS or oAβ, as in **(C**,**E)**. Two-way Anova, n=6-8 from 2 independent experiments. **(H)** Quantification of Homer1 engulfment in WT or *Spp1*^*KO/KO*^ P2Y12^+^ microglia, ICV treated with either PBS or oAβ, as in **(D**,**F)**. Two-way ANOVA, n=10-16 cells per brain from 2 independent experiments. ***p < 0.005 and ****p < 0.001. Data are shown as mean ± SEM.

### *Spp1* modulates complement activation and microglial synaptic engulfment upon acute oAβ challenge

Our data suggest that there is a significantly increased level of SPP1 protein within the brain parenchyma of pre-plaque *App*^NL-F^ hippocampi (Fig. 1), and that SPP1 is mainly produced by perivascular cells (Fig. 2). SPP1 induction coincides with the onset of microglia-synapse engulfment in the CA1 hippocampus. SPP1 has been associated with increased phagocytosis in the context of Aβ-challenge; however, the mechanisms by which SPP1 exerts these effects and whether it impacts microglia-synapse interaction are unknown^36^. Hence, we hypothesized that hippocampal perivascular SPP1 in the pre-plaque AD-like brains impacts phagocytic functions of neighboring parenchymal microglia, specifically their engulfment of synapses. To test this assumption, we used an *in vivo* model of oAβ-induced synapse engulfment (Fig. 3A)^12^, which allows to monitor microglia- and complement-mediated synapse engulfment in contralateral hippocampus of WT mice post intracerebroventricular (ICV) injection of oAβ [Aβ (1-40) S26C dimers]. We found significantly increased levels of SPP1 protein in the contralateral hippocampus at 18 h (but not at 72 h) after injection of oAβ versus PBS vehicle control, a time point when we observed microglia engulfing synapses (Fig. 3B). Upregulation of SPP1 also coincided with a significant upregulation and deposition of C1q protein (Fig. 3C,G). In line with C1q upregulation, hippocampal microglia of oAβ- injected animals displayed higher levels of Homer1-immunoreactive synapses inside CD68^+^ lysosomes, as compared to control mice (Fig. 3D,H).

To test whether SPP1 is required for microglial engulfment of synapses involving C1q, we performed ICV injections of oAβ or PBS in *Spp1*-deficient (*Spp1*^*KO/KO*^) mice^37^. Unlike in sex- and age-matched WT controls, oAβ failed to induce increase of microglial C1q in *Spp1*^*KO/KO*^ mice in the contralateral hippocampus (Fig. 3E,G). Further, microglia failed to increase engulfment of Homer1^+^ synapses in *Spp1*^*KO/KO*^ mice upon oAβ challenge (Fig. 3F,H). Together, these data demonstrate that SPP1 is necessary for microglia to upregulate C1q and engulf synapses in response to synaptotoxic Aβ. To test whether the failure for microglia to engulf synapses in *Spp1*^*KO/KO*^ mice was due to an intrinsic defective phagocytic capacity of microglia, we isolated primary microglia from WT and *Spp1*^*KO/KO*^ postnatal brains and assessed their *in vitro* ability to engulf oAβ-bound synaptosomes (Suppl Fig. 4A-D). We found that primary microglia from WT mice engulfed Aβ-bound synaptosomes and that this engulfment was similarly performed as well in primary microglia from *Spp1*^*KO/KO*^ mice (Suppl Fig. 4B). Of note, dose-response curve analysis revealed that primary microglial engulfment was stimulated by extracellular SPP1 (Suppl Fig. 4C). This establishes that the absence of microglia-synapse engulfment in the oAβ -challenged *Spp1*^*KO/KO*^ brain is not due to an intrinsic inability of *Spp1*^*KO/KO*^ microglia to phagocytose or respond to stimuli. Altogether, these results suggest that secreted SPP1, most likely from perivascular cells, in response to oAβ, facilitates microglial engulfment of synapses.

**Figure 4:**
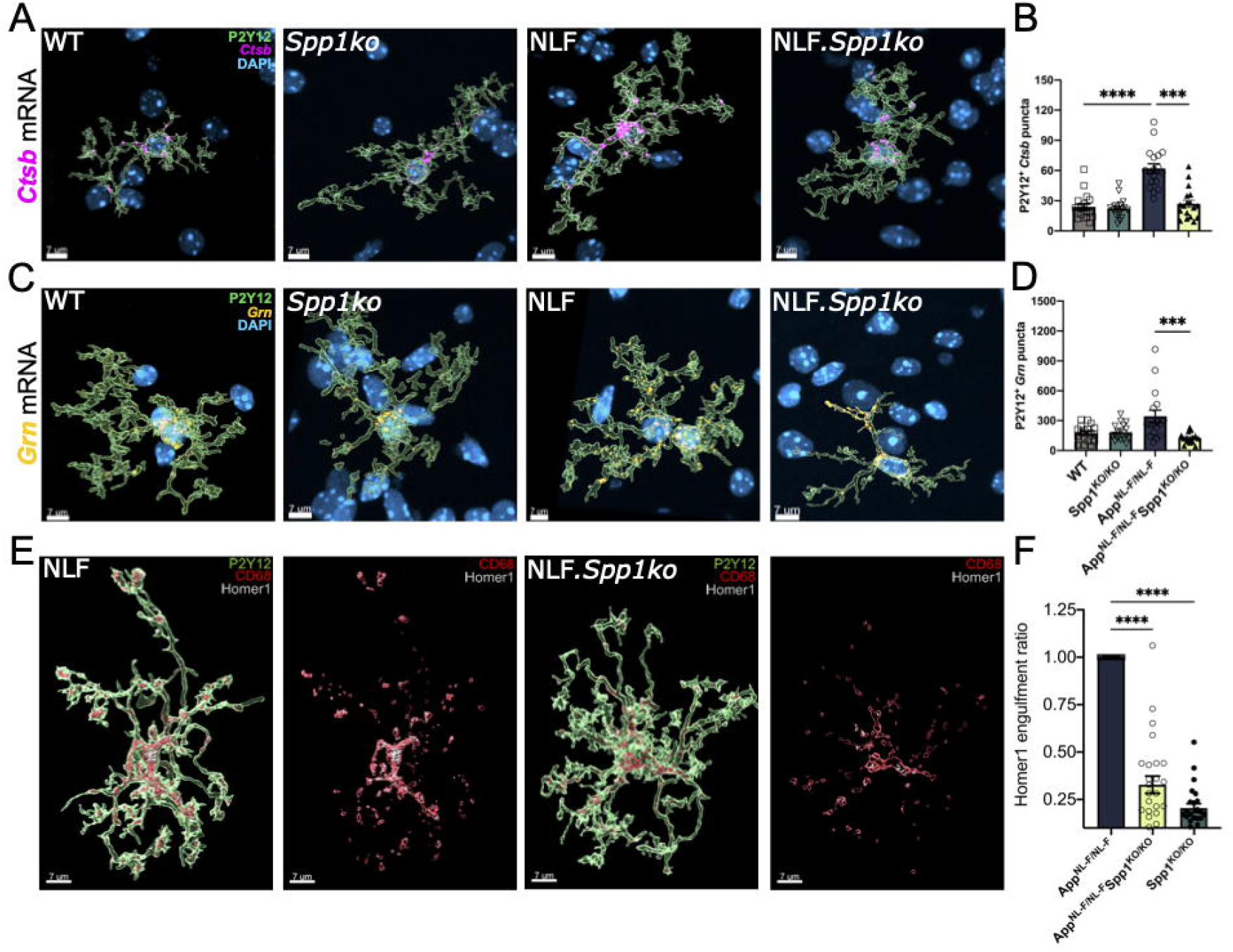
*Spp1* drives microglia phagocytic state and engulfment of synapses in pre-plaque *App*^NL- F^ hippocampus. **(A-D)** Representative image of 3D reconstructed P2Y12^+^ microglia expressing phagocytic markers *Ctsb* **(A**,**B)** and *Grn* **(C**,**D)** assessed by smFISH-IHC in 6 mo WT, *Spp1*^*KO/KO*^, *App*^NL-F^ and *App*^NL-F^.*Spp1*^*KO/KO*^ SLM. Quantification of *Ctsb* **(B)** and *Grn* **(D)** mRNA levels expression within microglia. One way ANOVA, Kruskal-Wallis n=6-9 cells per brain from n=3 mice for all genotypes. **(E)** Representative 3D reconstructed images showing Homer1 engulfment within CD68^+^ lysosomes of P2Y12^+^ microglia in 6 mo *App*^NL-F^ versus *App*^NL-F^.*Spp1*^*KO/KO*^ SLM. **(F)** Quantification of Homer1 engulfment ratio in P2Y12^+^ microglia of *App*^NL-F^ versus *App*^NL-F^.*Spp1*^*KO/KO*^ mice. One way ANOVA, Kruskal-Wallis, n=22 cells per brain from n=3 mice for all genotypes. Data are representative of 2 independent experiments and normalized to *App*^NL-F^ engulfment data. *p < 0.05 ***p < 0.005 and ****p < 0.001. Data are shown as Mean ± SEM.

### *Spp1* drives microglia phagocytic state and engulfment of synapses in pre-plaque *App*^NL-F^ hippocampus

We then asked whether SPP1 is required for the increased microglia-synapse engulfment in 6 mo *App*^NL-^ mice. To this end, we crossed *Spp1*^*KO/KO*^ mice with *App*^NL-F^ mice and analyzed them versus age- and sex-matched WT, *Spp1*^*KO/KO*^, and *App*^NL-F^ animals (Fig. 4). First, we assessed whether *Spp1* deficiency altered phagocytic signature in microglia and used smFISH-IHC to probe mRNA expression of lysosomal markers *Grn* (encoding for progranulin) and *Ctsb* (encoding for Cathepsin B) specifically in P2Y12^+^ microglia (Fig. 4A-D). *Grn* and *Ctsb* are key components of the endolysosomal processing machinery in microglia^10^. We found significantly decreased *Grn and Ctsb* mRNA expression in hippocampal microglia of 6 mo *App*^NL-F^. *Spp1*^*KO/KO*^ versus *App*^NL-F^ mice, as measured by mRNA puncta in 3D reconstructed P2Y12^+^ cells (Fig. 4A-D). In line, *Ctsb* mRNA expression was significantly increased in *App*^NL-F^ microglia compared to age-matched WT animals, while a trend towards increased *Grn* expressed was observed in microglia from *App*^NL-F^ mice, however without reaching significance (Fig. 4C,D). We next questioned whether increased microglial *Grn* and *Ctsb* expression coincided with dysregulated microglial–synapse interactions in 6 mo *App*^NL-F^ mice in the absence of *Spp1*. To assess this, we applied the synaptic engulfment assay to P2Y12^+^ microglia in CA1 hippocampus of *App*^NL-F^.*Spp1*^*KO/KO*^ mice compared to *App*^NL-F^ and *Spp1*^*KO/KO*^ as controls (Fig. 4E,F). We observed a significant drop in Homer1 immunoreactivity within microglial lysosomes of *App*^NL-F^.*Spp1*^*KO/KO*^ mice as compared to *App*^NL-F^ mice, suggesting that hippocampal microglia in pre-plaque burdened *App*^NL-F^ mice failed to effectively engulf synapses in the absence of SPP1 (Fig. 4E,F). We observed a further drop in microglial synaptic engulfment in *Spp1*^*KO/KO*^ as compared to *App*^NL-F^.*Spp1*^*KO/KO*^ mice, suggesting that additional mechanisms likely account for microglial engulfment of synapses (Fig. 4F). Together, our data suggest SPP1 mediates microglia-mediated synaptic engulfment in response to synaptotoxic Aβ and in pre-plaque *App*^NL-F^ mice.

### NicheNet reveals candidate perivascular-microglial interaction networks regulated by *Spp1*

Our results altogether suggest an SPP1-mediated crosstalk originating from the perivascular space to stimulate phagocytic capacity in microglia. To obtain insight into how SPP1 potentially regulates interactions between the perivascular cells, i.e., PVM and PVF, and parenchymal microglia in the hippocampus of *App*^NL-F^ mice, we performed comparative single cell RNA sequencing (scRNA-seq) analysis on sorted CD140a^+^ PVF, CD45^high^CD11^int^CD206^+^ PVM and CD45^int^CD11^int^CX3CR1^high^ microglia from dissected hippocampi of 6 mo *App*^NL-F^ mice versus *App*^NL-F^.*Spp1*^*KO/KO*^ animals (Suppl Fig. 5). Age-matched WT and *Spp1*^*KO/KO*^ mice were used as controls. We performed scRNA-seq using the 10X Genomics platform and after quality control, we ran unsupervised clustering and annotated cell types based on expression of known marker genes (Suppl Fig. 5A,B). Of note, the sorted CD140a^+^ cells also included oligodendrocyte precursors (OPCs), that share expression of CD140a but can be discerned according to expression of OPC-specific markers including *Lhfpl3, Sox6* and *Bcan* (Suppl Fig. 5B). Further, microglia expressed pan-markers such as *Sall1, Tmem119, P2ry12*, and PVF were positive for *Cdh5, Lama1* and *Dcn* that distinguished them from OPC. Finally, as previously described^5,38^, we confirmed *Mrc1, Pf4* and *Cd163* as specific markers enriched in PVM (Suppl Fig. 5B). Of note, in contrast to our observations using smFISH and super-resolution imaging methods, we did not detect *Spp1* mRNA in isolated PVM from either genotype. This conforms with the scRNA-seq dataset by Zeisel *et al* (2018) on multiple CNS cell types (Suppl Fig. 2D)^35^, although translatome analysis of PVM of brains of LPS endotoxin challenged *Cx3cr1*^ccre^ *Lyve1*^ncre^:RiboTag-mice has demonstrated the presence of *Spp1* in Lyve1^+^ PVM (Suppl Fig. 5C)^5,35^. This discrepancy of detected expression levels between scRNA-seq of isolated cells versus the RiboTag approach, which bypasses cell isolation and sorting^39,40^, and the highly sensitive smFISH methods suggests that *Spp1* signature within PVM is highly regulated and closely associated to changes to its microenvironment, as earlier demonstrated for many other microglial transcripts^3^. It further suggests that *Spp1*^+^ PVM likely do not survive isolation and highlights the relevance of studying SPP1 in an intact spatial context.

Next, we utilized NicheNet, which is a computational algorithm that predicts ligand-target links between interacting cells based on obtained transcript datasets and prior knowledge on molecular signaling and regulatory networks^41^. We applied NicheNet to investigate how intercellular communication between PVM/PVF and microglia is altered by *Spp1* deficiency, and how this affects microglial functional states in *App*^NL-F^ versus *App*^NL-F^.*Spp1*^*KO/KO*^ mice (Fig. 5). We considered SPP1 to function through either paracrine and/or autocrine mechanisms, i.e., modulating microglial function through direct or indirect signaling within the perivascular space. Among top predicted ligands affected in PVM and PVF from *App*^NL-F^.*Spp1*^*KO/KO*^ versus *App*^NL-F^ was transforming growth factor-beta 1 (TGF-β1), a cytokine critical for microglial development and previously shown to be modulated by SPP1 within fibroblasts (Fig. 5A-C)^42,43^. Other ligands include A Disintegrin and Metallopeptidase Domain 17 (ADAM17) and calreticulin, the latter being reported as a ‘eat-me signal’ driving macrophage phagocytosis of apoptotic cells^44^. Further, integrin receptors *Itgb5, Itgb1 and Itgav*, subunits of the SPP1 receptor, were affected in microglia of *App*^NL-F^.*Spp1*^*KO/KO*^ versus *App*^NL-F^ mice, likely reflecting *Spp1* deficiency. Notably, another affected receptor in microglia was *Tgfbr2* (Fig. 5D), a key regulator of microglial homeostasis (Fig. 5B-D)^45^. Finally, top affected target genes within microglia of *App*^NL-F^ mice deficient of *Spp1* include *Ccnd1* and protein tyrosine phosphatase *Ptpn6* (Fig. 5E), a major downstream signaling molecule of CD33 and upregulated in microglia surrounding Aβ plaques^46^. Altogether, NicheNet predicted both paracrine and autocrine crosstalk signals between the perivascular space and microglia in *App*^NL-F^ mice in the context of SPP1 signaling. Examples include the highly relevant TGF-β pathway and calreticulin (Fig. 5), thereby suggesting a role for perivascular SPP1 in modulating synaptic phagocytosis in microglia in AD mouse brains.

**Figure 5:**
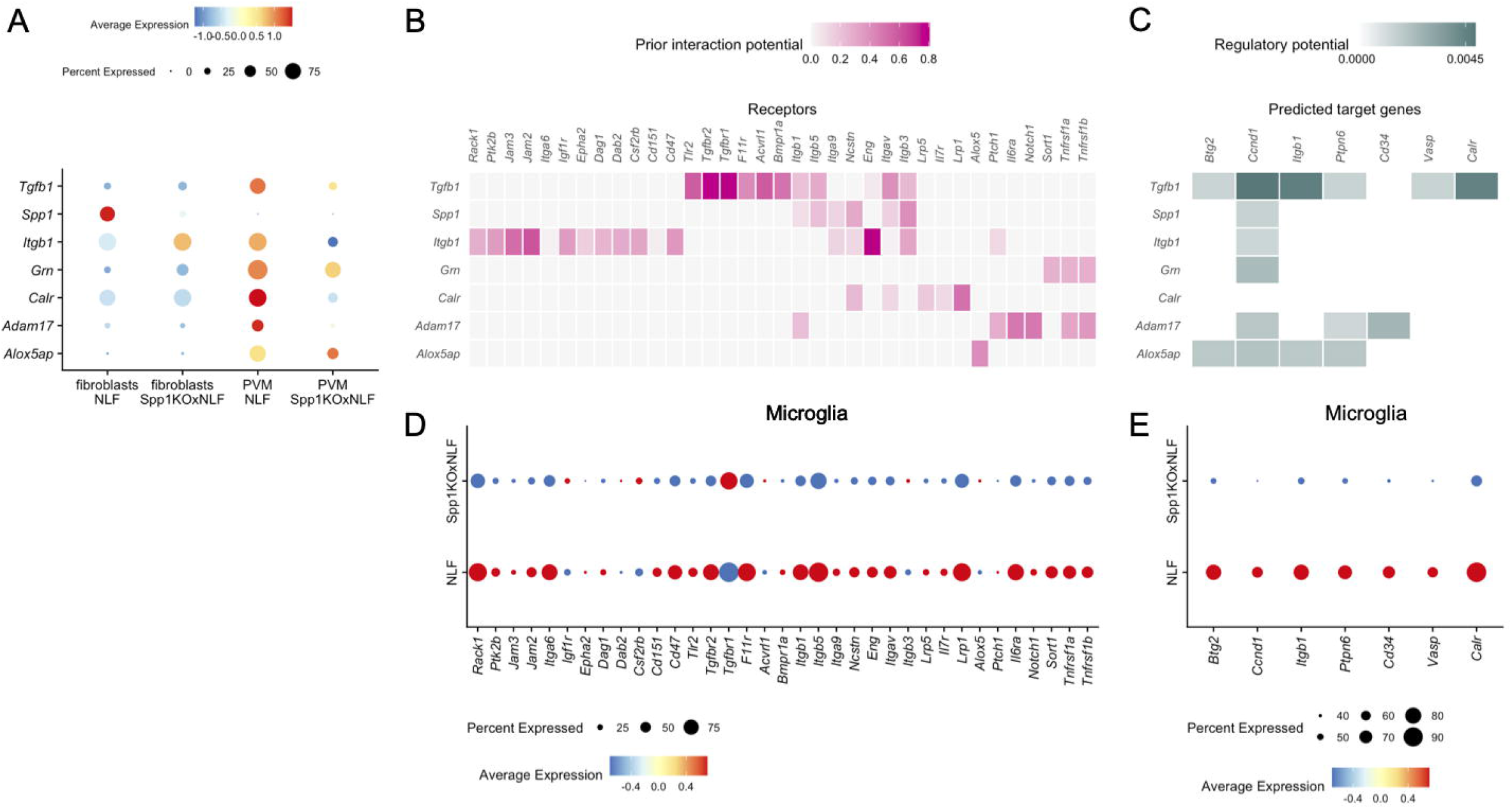
NicheNet reveals candidate perivascular-microglial interaction networks regulated by *Spp1*. **(A)** Expression level of selected ligands expressed by cell types known to express *Spp1* (PVF and PVM), by cell type and by genotype. Radius of dot is proportional to the percentage of cells expressing the gene, colour is the scaled gene expression level. **(B)** Predicted receptor genes for ligands represented in A, which show differential expression in microglia (receiver cells). Colour represents the predicted interaction potential. **(C)** Predicted target genes downstream of receptors identified in B, which show differential expression in microglia (receiver cells). Colour represents the predicted regulatory potential. **(D)** Expression of predicted receptor genes in microglia, by genotype. Radius of dot is proportional to the percentage of cells expressing the gene, colour is the scaled gene expression level. **(E)** Expression of predicted target genes in microglia, by genotype. Radius of dot is proportional to the percentage of cells expressing the gene, colour is the scaled gene expression level.

## Discussion

Genetic studies in sporadic AD highlight microglia and their role as professional phagocytes in disease pathogenesis. Microglia-synapse engulfment involving complement has been demonstrated to be relevant to synaptic loss and dysfunction in a wide variety of neurologic diseases across the lifespan, including neurodevelopmental and neuropsychiatric disorders, acute injury and virus-induced cognitive impairment, multiple sclerosis, and neurodegeneration^12,15,53,54,16,32,47–52^. Therefore, insight into mechanisms underlying microglia-synapse interaction will likely have a broad relevance to understanding region-specific synapse vulnerability in AD and other neurologic diseases involving synaptopathy.

Here we report perivascular SPP1 as a novel mediator of microglia-synapse engulfment in models of AD-like pathology. We found that in mice genetically deficient for *Spp1*, microglia fail to phagocytose synapses in the presence of early Aβ pathology (6 mo *App*^NL-F^) or upon acute challenge with synaptotoxic Aβ. SPP1 is a multifunctional glycoprotein that was originally identified as a proinflammatory cytokine secreted by T-cells and later found to be expressed in distinct tissue-resident macrophages linked with active clearance of apoptotic cells, chemotaxis and macrophage migration^20,55^. We demonstrate that SPP1 expression in the hippocampus of naïve unchallenged animals is restricted to PVM, border-associated macrophages at the perivascular space, and to the closely neighboring PVF, in WT mice. We corroborate this finding by colocalizing *Spp1* mRNA using pan-PVM markers including *Pf4, Cd163* and CD206 and also a CLEM study in *Spp1*^TdT^ mice confirmed PVM as primary SPP1^+^ cells in the hippocampus. The perivascular localization of SPP1 was recapitulated in post-mortem human brains of AD patients. In contrast, we did not observe any appreciable levels of microglial *Spp1* in the hippocampus of *App*^NL-F^ mice until they are plaque-burdened at 15 mo, likely reflecting disease-associated microglia that appear and cluster around plaques^25^. Indeed, SPP1 appears to be spatiotemporally regulated in a cell-type specific manner, depending on context, age, and brain region. We found PVM as a cellular origin for SPP1 in early *App* models when oAβ deposits along the vasculature, consistent with a recent report that suggested *Spp1* upregulation in Lyve1^+^ PVM upon LPS challenge in adult mice^56^. Conversely, in the perinatal and prenatal brain, *Spp1* is expressed by microglia and associates with axon tracts of the *corpus callosum*^5,23,57^. Of note, glutaminergic and GABAergic neurons in adult hindbrain are also shown to express SPP1^35^. Although gross levels of BBB impairment and infiltration of immune cells into the brain parenchyma are thought to occur in later stages of amyloidosis^58^, we currently cannot fully exclude the contribution of SPP1 from circulating lymphocytes. Future studies allowing cell-type specific fate mapping and mutagenesis of hippocampal PVM and PVF subsets will be needed to provide insight into the origin and fate of SPP1-expressing perivascular subsets.

What could be the relevance of elevated SPP1 originating from the perivascular space? In AD patients, SPP1 is found to be elevated in CSF^27–29^, which flows via the perivascular space towards the subarachnoid space. PVM are distributed along the perivascular space where they may play a critical role as sentinels and contribute to neurovascular and cognitive dysfunction in mouse models of amyloidosis. Exact mechanisms yet have been unclear^59^. Clearance of Aβ from brain parenchyma across the BBB represents an important homeostatic function, the impairment of which was linked to exacerbated vascular as well as parenchymal deposition of Aβ^*60,61*^. Blood vessels therefore may represent a frequent, early and vulnerable site for Aβ deposition; indeed it was postulated that early failure of perivascular drainage could further contribute to parenchymal Aβ deposition and associated neuronal toxicity^62^. Our findings of oAβ deposition along the vasculature in early stages of amyloidosis in *App*^NL-F^ mice support this view. This is further corroborated by the commonly observed vascular pathology in AD, afflicting approximately 80 % of patients^63^. Of note, an earlier study suggested a role for SPP1 in clearance of vascular Aβ deposits in transgenic APP/PS1 mice^36^. Altogether, these published reports and our data suggest a possible upregulation of SPP1 by PVM and PVF in response to Aβ deposition in the perivascular space.

Our observation that SPP1 signaling promotes synaptic engulfment by microglia in the hippocampus of AD models suggests that SPP1 functions to regulate phagocytic cell states. Indeed, in the absence of *Spp1*, microglia fail to upregulate key phagocytic genes such as progranulin and *Ctsb* in 6 mo *App*^NL-F^ mice. Of note, phagocytosis induced by microglial progranulin as well as complement proteins including C3 has been considered protective against Aβ plaque load and neuronal loss in plaque-rich mouse models of AD. In line with our data, this suggests that the timing of phagocytic activity in different stages of disease might be critical to determine the protective versus detrimental nature of the outcome^12,17,64^. Another interesting question raised is how SPP1 modulates microglial phagocytosis in the presence of Aβ pathology. SPP1 functions as a secreted or intracellular isoform^65^. Based on our data, a direct effect of intracellular SPP1 on microglia function seems unlikely in our experimental setting. First, we found that microglia-intrinsic *Spp1* signaling is not required for the engulfment of Aβ-treated synaptosomes, where primary microglia derived from WT and *Spp1*^*KO/KO*^ mice exhibit similar engulfment capacities of synaptosomes in the presence of extracellular SPP1. This is consistent with work in primary gut macrophages from *Spp1*-deficient mice, whereby low doses of recombinant SPP1 restored phagocytic function towards opsonized bacteria, akin to what we observed for primary microglia and synaptosomes^66^. Second, we did not observe mRNA or protein SPP1 expression localized inside hippocampal microglia at the age of 6 mo, yet the same cells fail to engulf synapses in *App*^NL-F^.*Spp1*^*KO/KO*^ mice or *Spp1*^*KO/KO*^ mice challenged with synaptotoxic Aβ. Third, we found increased protein levels of SPP1-immunoreactive puncta within the SLM parenchyma using super-resolution imaging, suggesting that the observed punctate signals are mostly extracellular. Altogether, these results suggest that SPP1 which is secreted by PVM and PVF influences microglia phagocytosis.

Precise mechanisms of how microglial engulfment is initiated by SPP1 are to be investigated. SPP1 could opsonize neuronal material e.g. damaged synapses for microglial phagocytosis^67^. It could also engage directly with the canonical receptors, α_v_β_3_, which are expressed on microglia, to trigger downstream signaling pathways. Indeed, using NicheNet, we see *Itgb3* and *Itgav*, which encodes α_v_β_3_, to be downregulated in microglia of *App*^NL-F^.*Spp1*^*KO/KO*^ mice. Our NicheNet analysis further shows reduced expression levels of both *Tgfb1* in PVM and its receptor *Tgfbr2* in microglia from *App*^NL-F^.*Spp1*^*KO/KO*^ mice, suggesting that SPP1 potentially coordinates PVM-microglial crosstalk via autocrine TGF-β signaling. TGF-β is a major determinant of microglia maturation and homeostasis, and *Tgfbr2* deficient microglia demonstrate dysregulated expression of transcripts related to phagosome formation and immune activation^43,45^. Interestingly, TGF-β1 has been shown to be increased in CSF and perivascular space of AD patients, where its levels positively correlate with Aβ deposition along blood vessels, and TGF-β1- overexpressing mice show enhanced microglial phagocytosis and plaque amelioration in a complement-dependent manner^68,69^. Other predicted *Spp1*-specific pathways via NicheNet in microglia of *App*^NL-F^ mice include *Calr*, encoding for calreticulin, a multifunctional chaperone protein that has been described as an ‘eat-me’ signal^44^. Altogether, our results suggest multiple mechanisms by which microglial phagocytosis could be modulated by perivascular SPP1 signaling. Determining molecular mechanisms of how these pathways may contribute to synaptic engulfment will be important to better understand multi-cellular events and crosstalk between perivascular space and the brain parenchyma.

Overall, our study suggests that cellular and molecular crosstalk between the perivascular space and microglia via SPP1 mediates synaptic engulfment in pre-plaque AD. In peripheral tissues, insight into stroma-macrophage interactions demonstrates intricate and functionally relevant immune crosstalk that underlies critical tissue remodeling and homeostasis. Likewise, our results highlight a potential role for perivascular-microglia interactions in the brain. As such, the presence of a SPP1-producing perivascular niche may offer opportunities to specifically manipulate and block microglia-mediated synaptic engulfment.

## Acknowledgements

We thank Mari Shinohara (Duke) for valuable discussion and input. We thank Steve West (Sainsbury Wellcome Centre, UCL), Isabelle Noelle Chiong (King’s College London), Emir Turkes (UK DRI at UCL), Javier Rueda-Carrasco (UK DRI at UCL) and Phillip Muza (UCL) for experimental help. We thank Frances Edwards (UCL; PPL); Takaomi Saido (Riken; *App*^NL-F^ mice), Elena Ghiradello and Phillip Muckett (UK DRI at UCL; animal husbandry). We thank Virginia Lee (UPenn; NAB61 antibody), John Cirrito (WUSTL; mHJ3.4antibody) and Nic Cade (UK DRI at UCL for microscope imaging assistance). We thank Jamie Evans (FACS Core Facility at UCL, flow cytometry assistance and cell sort). We would like to acknowledge The Jackson Laboratory’s Genetic Engineering Technologies Scientific Service for mouse strain development. This work was supported by the UK Dementia Research Institute (SH) (UKDRI-1011) (which receives its funding from UK DRI Ltd, funded by the UK Medical Research Council, Alzheimer’s Society and Alzheimer’s Research UK), Bright Focus Foundation Grant (SH) (183609) and UCL Neurogenetic Therapies Programme funded by Sigrid Rausing Trust. SDS is supported by a Wellcome Trust Sir Henry Wellcome Fellowship (221634/Z/20/Z), GC by a Wellcome Trust 4-year Neuroscience PhD studentship (219906/Z/19/Z). The Queen Square Brain Bank is supported by the Reta Lila Weston Institute of Neurological Studies, UCL Queen Square Institute of Neurology. JJB is funded by core funding to the MRC Laboratory for Molecular Cell Biology at University College London (award code MC_U12266B). The LMCB volume EM facility was set up with funding from the Wellcome Trust (218278/Z/19/Z). The *Spp1*^TdT^ reporter mouse model was developed with funding from an anonymous organization. The authors would like to acknowledge helpful discussions and feedback from the International Neuroimmune Consortium.

## Author Contributions

Conceptualization, S.D.S., S.H., S.J.; Methodology, S.D.S., S.H., D.G., M.S., C.S.F., J.J.B.; Software, J.J.B. and C.S.F.; Validation, S.D.S., J.Z.G., G.C., L.S.S.F., C.T., D.S., T.C., T.L., J.J.B., M.S.; Formal Analysis, S.D.S., J.Z.G., L.S.S.F., D.G., T.C., J.J.B., C.S.F.; Investigation, S.D.S, J.Z.G., G.C., L.S.S.F., C.T., D.S., T.C., T.L., J.J.B., M.S., C.S.F.; Resources, D.G., C.T., T.L., J.J.B., M.S., C.S.F.; Data Curation, C.S.F.; Writing – Original Draft, S.D.S and S.H.; Visualization, S.D.S., J.J.B., C.S.F.; Supervision, S.H.; Funding Acquisition, S.H.

## Lead Contact

Soyon Hong

## Star Methods

### Mice

Experiments were performed in accordance with the UK Animal (Scientific Procedures) Act, 1986 and following local ethical advice. Experimental procedures were approved by the UK Home Office and ethical approval was granted through consultation with veterinary staff at University College London.

For all experiments, C57BL/6J (WT) mice were obtained from Charles River UK. *Spp1*^*KO/KO*^ (B6.129S6(Cg)-Spp1^tm1Blh^/J; Stock no. 4936), and CX_3_CR-1^GFP^ (B6.129P2(Cg)-Cx3cr1^tm1Litt^/J; Stock no. 5582) mice were obtained from The Jackson Laboratory (Bar Harbor, Maine, USA). *App*^NL-F/NL-F^ mice were obtained from Prof Frances Edwards (UCL, London, UK)^31^.

All animals were housed under temperature-controlled pathogen-free conditions with 12 h light/dark cycle with ad libitum supply of food and water. Both male and female age-matched mice were used in this study.

### *Spp1* reporter mouse model development

All animal work was approved by the Jackson Laboratory Animal Care and Use Committee and adhered to the standards of Guide for the Care and Use of Laboratory Animals set forth by the NIH.The *Spp1*^*tm1(tdTomato)Msasn*^ mouse allele was generated using direct delivery of CRISPR-Cas9 reagents to mouse zygotes. An IRES-tdTomato construct was introduced in the mouse Spp1 gene (Ensembl Gene UID ENSMUSG00000029304) (Suppl Fig. 3A). Analysis of genomic DNA sequence surrounding the target region, using the Benchling (www.benchling.com) guide RNA design tool, identified a gRNA sequence (AACAAGAAAAAGTGTTAGTG) with a suitable target endonuclease site at the stop codon of exon 7 of the mouse *Spp1* locus. *Streptococcus pyogenes* Cas9 (SpCas9) V3 protein and gRNA were purchased as part of the Alt-R CRISPR-Cas9 system using the crRNA:tracrRNA duplex format as the gRNA species (IDT, USA). Alt-R CRISPR-Cas9 crRNAs (Product# 1072532, IDT, USA) were synthesized using the gRNA sequences specified in the DESIGN section and hybridized with the Alt-R tracrRNA (Product# 1072534, IDT, USA) as per manufacturer’s instructions. A plasmid construct with a 2kB 5’ homology arm ending 62bp past the Spp1 stop codon in exon 7, IRES, tdTomato coding sequence, bGH poly(A) signal and 1.5kB 3’ homology arm was synthesized by Genscript (Suppl Fig. 3B). To prepare the gene editing reagent for electroporation, SpCas9:gRNA Ribonucleoprotein (RNP) complexes were formed by incubating AltR-SpCas9 V3 (Product#1081059, IDT, USA) and gRNA duplexes for 20 min at room temperature (RT) in embryo tested TE buffer (pH 7.5). The SpCas9 protein and gRNA duplex were at 833 ng/*μ*l and 389 ng/ *μ*l, respectively, during complex formation. Post RNP formation, the purified plasmid was added and the mixture spun at 14K RPM in a microcentrifuge. The supernatant was transferred to a clean tube and stored on ice until use in the embryo electroporation procedure. The final concentration of the gRNA, SpCas9 and plasmid components in the electroporation mixture were 600 ng/*μ*l, 500 ng/*μ*l and 20 ng/*μ*l, respectively. Fertilized mouse embryos were generated via natural mating and cultured as described previously^70^. C57BL/6J (Stock# 000664, The Jackson Laboratory, USA) donor female mice (3–4 weeks of age) were superovulated by administration of 5 IU of pregnant mare serum gonadotrophin (PMSG) via intraperitoneal (ip) injection (Product# HOR-272 ProSpec, Israel) followed 47 h later by 5 IU (ip) human chorionic gonadotrophin (hCG) (Product# HOR-250, ProSpec, Israel). Immediately postadministration of hCG, the female was mated 1:1 with a C57BL/6J stud male and 22 h later checked for the presence of a copulation plug. Female mice displaying a copulation plug were sacrificed, the oviducts excised, and embryos collected. Electroporation was performed as described^70^. In brief, zygotes were treated with the acidic Tyrode’s solution (Product# T1788, Millapore-Sigma, USA) for 10 s and washed extensively in pre-warmed M2 media (Product# M7167, Millapore-Sigma, USA). Zygotes were then placed in 10 *μ*L drops of Opti-MEM media (Product# 3198570, ThermoFisher-Gibco, USA). 10 *μ*L of gene editing reagent solution, including the SpCas9/gRNA and the ssDO, was mixed with the Opti-MEM drops with the embryos and deposited into a 1 mm electroporation cuvette (Product# 45-0124, Harvard Apparatus, USA). Electroporation was performed in an ECM830 Square Wave Electroporation System (Product# 45-0661, BTX, USA). The electroporation setting was a 1 ms pulse duration and two pulses with 100 ms pulse interval at 30 V. Following the electroporation, a pre-warmed 100 *μ*L aliquot of M2 media was deposited into the cuvette with a sterile plastic pipette to recover the embryos. The zygotes were removed from the cuvette and washed in pre-warmed M2 media. Embryos were immediately transferred into B6Qsi5F1 pseudopregnant female mice, a F1 hybrid strain produced by breeding C57BL/6J female mice with the inbred Quackenbush Swiss line 5 (QSi5) mouse strain^71^. Founders were first assayed by short range PCR with primer SR-FF 5’-TAATAATGGTGAGCAAGGGCGA-3’ and reverse primer SR-RR 5’-CTTTGATGACGGCCATGTTGTT-3’ within the reporter construct (see schematic). Positive founders were then screened by PCR: across the 5’ homology arm with primer 5’LR_F 5’-GAAAGTGCCTACTCGTGCCT-3’ and reverse primer 5’LR_R: CACATTGCCAAAAGACGGCA-3’; across the 3’ homology arm with primers 8431 GCATCGCATTGTCTGAGTAGGT and 3’LR_R: ccatcatggctttgcatgac; and for the presence of plasmid backbone with primers 8431: GCATCGCATTGTCTGAGTAGGT and 9581: AGCGCAACGCAATTAATGTG. Sanger sequencing was performed across the homology arm junctions and portions of the knocked-in reporter gene (Suppl Fig. 3C).

Founders were selected that: were positive by short-range PCR assays; had appropriate sequence across the homology arm junctions; were negative for the plasmid backbone; and had correct sequence of the inserted construct. These were bred to C57BL/6J.

Once the line was established, mice were genotyped using a Taqman qPCR protocol run on a real time PCR instrument (Roche LightCycler480). Forward primers for the wild-type allele (AAACACAGTTCCTTACTTTGCAT producing a 94bp product) and the mutant allele (AGGATTGGGAAGACAATAGCA in the bGH poly A producing a 84bp product) were combined with a common reverse primer (CACTGAACTGAGAAATGAGCAGT) using an annealing temperature of 60°C. The wild-type probe (5’ HEX fluorophore label) used was TGTTAGTGAGGGTTAAGCAGGAATA and the knock-in probe (5’ FAM fluorophore label) used was ATGCGGTGGGCTCTATGG, each utilizing a Black Hole Quencher (BHQ) on their 3’ end. These were run with an EndPoint protocol, imaging negative (quenched) fluorescent values at the completion of the cycling protocol.

This novel line of *Spp1*-IRES-TdTomato mice is available as B6J.*Spp1*^*tm1(tdTomato)Msasn*^/J (Stock no. 33731) from The Jackson Laboratory (Bar Harbor, Maine, USA).

### Intracerebroventricular injections of S26C oAβ

S26C oAβ dimers were purchased from Phoenix Pharmaceuticals (Burlingame, CA, USA)^12^. Adult (2-3 mo) mice were administered 4 % isoflurane in oxygen for anesthesia induction followed by maintenance at 1.5–2 % isoflurane during surgery. Surgery was performed after head fixing in a stereotaxic frame (World Precision Instruments). Sterile eye drops were applied to mouse eyes during surgery to prevent drying (Viscotears, Bausch and Lomb). Marcaine (0.025 %) was applied locally, the skull was exposed by a single incision along the midline and a unilateral craniotomy was drilled with a 0.9-bit drill burr (Hager and Meisinger). Pulled long-shaft borosilicate pipettes (Drummond Scientific) were backfilled with mineral oil before loading with oAβ (1 ng/µL) or sterile PBS vehicle. 4 µL total volume was injected into the right lateral ventricle (stereotaxic coordinates in millimeters from Paxinos and Franklin’s The Mouse Brain in Stereotaxic Coordinates, Fourth Edition; AP: -0.40, ML: 1.00, DV: -2.50) using a Nanofil 10 mL syringe (World Precision Instruments) connected to an UltraMicroPump-3 (World Precision Instruments) at a flow rate of 400 nL/min. The pipette was left in place for 5 min after complete substance injection and slowly withdrawn to avoid backflow along the pipette track. The incision on the scalp was closed with Vetbond tissue adhesive (3M). Subcutaneous carprofen (Carprieve, 5 mg/g body weight) and buprenorphine (Vetergesic, 0.1 mg/g body weight) diluted in 0.9 % saline were administered peri-operatively. Animals received carprofen (33.33 mg/mL) in their drinking water until tissue harvesting. The left hemisphere, contralateral to the injection site, was analyzed.

### Immunohistochemistry

Mice were deeply anesthetized and toe pinch tested before transcardial perfusion was performed by infusing 25-30 mL of filtered PBS, followed by 20 mL of 4 % ice-cold ultra-pure PFA (Generon 18814-20, Slough, UK). Brains were removed from the skull and fixed in 4 % PFA (Generon) for 24 h at 4 °C. For cryoprotection, brains were rinsed in PBS for at least 2 h to remove excess PFA and placed in 30% sucrose (in PBS). After 48 h, brains were transferred into tissue embedding molds (Generon 18646A-1, Slough, UK) filled with OCT (Sakura, Tokyo, Japan) and incubated at RT for 1 h followed by incubation in a sealed box of dry ice to be frozen and stored at -80 °C freezer. For cryosectioning, tissue blocks were removed from -80°C and placed in the cryostat microtome chamber (Leica CM1860 UV) for at least 30 min. The brains were sectioned either coronally or sagittally. 15 µm tissue sections were collected onto Superfrost Plus GOLD slides (K5800AMNZ72, Thermo Scientific, Waltham, MA) or 30 µm sections were collected into cryoprotect solution in 24-well plates. Sections were blocked with 5 % BSA, 0.2 % Triton X-100 (Sigma-Aldrich 9002-93-1, Darmstadt, Germany) and 5 % Donkey Serum (Abcam ab7475, Cambridge, UK) in PBS at RT for 90 min before incubation with primary antibodies at 4 °C overnight. All primary antibodies were diluted in blocking buffer. After 4X PBS washes for 15 min each, sections were incubated with secondary antibodies in blocking buffer for 2 h at RT and then washed again with PBS. All secondary antibody aliquots were centrifugated at 15000 rpm for 15 min before being diluted in blocking buffer. The sections were then incubated for 5 min in DAPI (1/2000, Thermo Fisher Scientific D1306, Waltham, MA, USA) diluted in PBS before being mounted either in Prolong Gold Antifade Mounting Medium (Thermo Fisher Scientific P36930) for confocal imaging (Carl Zeiss) or in Prolong Glass Antifade Mountant (P36982) for STED imaging (Leica Microsystems).

For immunostaining of C1q and synapses, 30 µm free-floating tissue sections were washed in PBS on the perturbator followed by pre-treatment in 1 % Triton X-100 in PBS for 20 min and a quick rinse in PBS. Sections were then blocked in 20 % NGS, 1 % BSA, and 0.3 % Triton, in PBS for 2 h followed by standard primary antibody incubation overnight at 4 °C. Sections are washed in 0.3 % Triton X-100 in PBS for 30 min followed by secondary incubation for 4 h at RT and wash for 30 min. Sections were then incubated in 1:10000 DAPI in PBS with 0.3 % TX for 10 min and washed for 15 min. Finally, sections were mounted onto slides with prolong gold mounting medium and procured for at least 24 h before imaging.

### Post-mortem human brain tissue

Brains were donated to the Queen Square Brain Bank (QSBB) for neurological disorders (UCL Queen Square Institute of Neurology). All tissue samples were donated with the full, informed consent. Accompanying clinical and demographic data of all cases used in this study were stored electronically in compliance with the 1998 data protection act and are summarized in Suppl. Table 1. Ethical approval for the study was obtained from the NHS research ethics committee (NEC) and in accordance with the human tissue authority’s (HTA’s) code of practice and standards under license number 12198, with an approved material transfer agreement. The cohort included pathologically diagnosed cases of AD (n=3) and neurologically normal controls (n = 3). The level of AD pathology in all cases were assessed using current diagnostic consensus criteria^72,73^. The *APOE* genotype was also determined for each case as previously described^74^.

### Immunohistochemistry on post-mortem human brain

Slides with 8 µm mounted tissue sections from the frontal cortex were incubated at 60 °C overnight. Sections were deparaffinized in Xylene and rehydrated in decreasing grades of alcohol. Slides were incubated in methanol/hydrogen peroxide (0.3%) solution for 10 min to block endogenous peroxidase activity. For heat-induced antigen retrieval, slides were then transferred to a boiling solution of 0.1 M citrate buffer (pH 6.0) and pressure cooked at maximum pressure for 10 min. Non-specific binding was blocked by incubating slides in 10 % non-fat milk for 30 min at RT. Sections were incubated in anti-SPP1 antibody for 1 h at RT. After three gentle 5 min washes in tris-buffered saline with tween (TBS-T); slides were incubated for 45 min in biotinylated goat anti-mouse IgG secondary antibody (Vector Laboratories BA 9200, 1:200). Slides were washed and incubated in Avidin-Biotin Complex (ABC; Vector) for signal amplification. The slides were then washed for a final time and 3,3’-Diaminobenzidine (DAB) used as the chromogen and counterstained in Mayer’s haematoxylin (BDH). Finally, slides were dehydrated in increasing grades of alcohol (70, 90 and 100 % IMS), cleared in xylene and mounted.

Double immunofluorescence staining was carried out on the frontal cortex. Sections were prepared as detailed above up to the incubation of the anti-SPP1 antibody, secondary biotinylated goat anti-mouse and ABC. Antibody binding was visualized using a TSA Cyanine 3 amplification kit (Perkin-Elmer) which was applied to sections for 20 min at RT. After TBS-T washing, sections were incubated with anti-CD206 antibody for 1 h at RT and species-appropriate Alexa Fluor 658 secondary antibodies (Invitrogen, 1:1000) for 2 h at RT to visualise the antibody. Sections were washed a final three times in TBS-T with the second wash incorporating a 10 min incubation with 4’,6-diamino-2-phenylindole (DAPI, Invitrogen, 1:1000) nuclei counterstain. Slides were mounted using Vectashield anti-fade mounting medium (Vector Laboratories)

### Image Acquisition

Images were acquired using a Zeiss LSM800 confocal microscope (40x objective, 1.3NA oil, 20x objective 0.8NA, 63x 0.8NA oil). Settings were kept constant for all sections in the same comparison group. Step size was determined using the optimal interval adjustment on the Zen blue software with a stack size of 10-14 µm for all on-slide IHC experiments and 6-10 µm for RNAScope experiments.

### Secreted SPP1 fluorescence intensity analysis

For quantification of secreted SPP1, Triton X-100 was excluded from the blocking buffer to retain signals of secreted and membrane-bound protein immune-reactivity. To quantify SPP1 protein expression, images were processed in Fiji ImageJ (NIH, Bethesda, USA)^75^. An automated ImageJ macro was created to analyse the signal intensity of each slice in the z-stack and select the plane with the highest signal intensity. Background was subtracted with a rolling ball radius of 10. Images were thresholded with consistent thresholding parameters and made into binary images based on intensity. As pixel intensity information has been translated into area, particle analysis function was used to quantify the total immune-reactive area.

### Antibodies

For immunostaining, all the antibodies used were rabbit anti-mouse C1q (1/200, Abcam, Cambridge, UK), goat anti-mouse SPP1 (1/50, Bio-Techne, Minneapolis, MN, USA), rabbit anti-mouse IBA1 (1/500, Wako Chemicals, Osaka, Japan), rabbit anti-mouse GLUT1 (1/10000, Merck Millipore, Burlington, MA, USA), rat anti-mouse CD68 (1/200, Bio-rad, Hercules, CA, USA), rabbit anti-mouse P2Y12 (1/500, Anaspec, Fremont, CA, USA), rat anti-mouse CD206 (1/200, Bio-rad), NAB61(kindly provided by Virginia M-Y Lee, Philadelphia), HJ5.1 (kindly provided by John R. Cirrito, St. Louis), Homer1 (160006, Synaptic Systems), 6E10 (1/200, BioLegend), 4G8 (1/200, BioLegend), and CD140a (1/50, Cell Signaling Technology). Secondary antibodies used were a combination of Alexa Fluor 488, 594, and 647 (1/200, Jackson ImmunoResearch and Fisher Scientific) chosen from goat anti-rabbit, goat anti-rat, goat anti-mouse, donkey anti-goat, donkey anti-rat, donkey anti-mouse and donkey anti-rabbit secondaries. For flow cytometry, BUV395 CD45 (BD Biosciences), Pe-Cy7 CD11b (BD Biosciences), BV421 CX3CR1 (Biolegend), PE CD140a (Miltenyi), APC CD206 (Biolegend) were used.

### *In vivo* microglial engulfment analysis

Engulfment analysis was performed as previously described^12^. 30 µm free-floating tissue sections were immunostained with IBA1, CD68 and Homer1. For each mouse, 4-6 regions of interest within CA1 SLM were acquired using a 63X 1.4-NA objective on a Zeiss 800 microscope. Next, 60-80 z-stack planes were taken with 0.27 *μ*m spacing and raw images were processed in Imaris (Bitplane) for analysis, after background subtraction of Homer1 channel. P2Y12^+^ cells and CD68 lysosomes were surface rendered with 0.25 *μ*m and 0.1 *μ*m smoothing, respectively. A mask was applied within P2Y12^+^ CD68^+^ reconstructed lysosomes for Homer1, and percentage engulfment of Homer1 within lysosomes was calculated using following formula: Volume of engulfed material (Homer1 within CD68) / Total microglial volume X 100.

### *smFISH* (RNAscope) and *smFISH* combined with IHC (smFISH-IHC)

To detect single RNA molecules, the RNA probes for *Spp1*(435191), *C1q*(441221), *Bin1*(529541), *Ctsb*(561541), *Grn*(422861), *Cd163*(406631), *Pdgfra*(480661), *Pf4*(502391) and *Tmem119*(472901) were purchased from Advanced Cell Diagnostics. RNAscope was performed following the manufacturer`s protocol in RNAscope®Fluorescent Multiplex Assay (320293). 15 *μ*m frozen brain sections were collected on Superfrost Plus GOLD Slides (Thermo Fisher, K5800AMNZ72) and dried overnight at 40 °C. Briefly, slides were incubated in H_2_O_2_ for 4 min at RT and washed in RNAse free water. Slides were placed in boiling Target Antigen Retrieval for 4 min and dehydrated in ethanol 100 % for 5 min. After air drying RT, the brains were treated with Protease Plus for 15 min at RT and incubated with the probes at 40 °C for 2 h. Further protocol as described by the manufacturer. For post-RNAscope immunostaining, brain sections were blocked in 5% normal goat serum (ab7481, Abcam, Cambridge, UK) with 1 % BSA and 0.1 % Triton x-100 in PBS at RT for 1 h followed by incubation in primary antibodies at 4 °C overnight and standard secondary staining procedure. RNAscope and immunofluorescent slides were acquired at*e*63X magnification and analyzed using ImageJ (NIH) and/or Imaris (Bitplane). For quantify single *Spp1* mRNA molecules, the “spots” tool in Imaris was used. First, microglia were 3D reconstructed using the same steps as described earlier. The channel marking the *Spp1* was masked by the surface of selected microglia. Using the quality filter, the diameter and number of spots were determined accordingly to the fluorescence signal. RNA expression was quantified based on the number of spots modelled in IMARIS.

### RNA isolation, reverse transcription and RT-qPCR

Mice were anaesthetized and perfused with ice-cold PBS. Next, fresh cortex and hippocampus were dissected and homogenized with TissueLyser II (QIAGEN) using the mRNA easy mini kit (QIAGEN) as described by the manufacturer. Next, RNA purity and concentration was assessed by Nanodrop. mRNA was converted to cDNA using the qScript cDNA SuperMix reverse transcription kit as described by the manufacturer (95048, Quantabio). For RT-qPCR, 12 ng of cDNA was loaded in triplicates per gene in a total volume of 20 *μ*l using the SYBR green PCR master mix as described by the manufacturer (4309155, ThermoFisher). The reaction was run using a LightCycler 96 Instrument (Roche) with white 96-well plates (04729692001, Roche). Triplicate Ct values were averaged and data is shown as respective to the geomean of 3 housekeeping genes (*Actb, Gapdh, Rpl32*) using the Ct delta method (2–*ΔΔ*Ct). Primers purchased from IDT were used at a concentration of 200 nM. *Actb*, Forward: CATTGCTGACAGGATGCAGAAGG, Reverse: TGCTGGAAGGTGGACAGTGAGG; Gapdh, Forward: CATCACTGCCACCCAGAAGACTG,Reverse: ATGCCAGTGAGCTTCCCGTTCAG; *Rpl32*, Forward: ATCAGGCACCAGTCAGACCGAT, Reverse: GTTGCTCCCATAACCGATGTTGG. *Spp1*, Forward: ATC TCACCATTCGGATGAGTCT, Reverse TGTAGGGACGATTGGAGTGAAA

### Microglia isolation for flow cytometry and FACS sort

Mice were deeply anesthetized and toe pinch tested before perfusion. After transcardial perfusion with 25-30 mL of filtered PBS, brains were quickly isolated from skull and hippocampus was dissected on ice using chilled instruments. For scRNA sequencing, brains were perfused with inhibitor cocktail including Actinomycin D (5 *μ*g/mL) and triptolide (10 *μ*M). Next, single cell suspension was prepared using the Adult Brain Dissociation kit from Miltenyi Biotec (Bergisch Gladbach, Germany), according manufacturer’s instructions and ^26^. Briefly, chopped tissue was incubated for 30 min in a mix of buffer Z with enzymes P, A and Y prepared according to the manufacturer’s instructions. Mechanical dissociation steps were performed at 10 min intervals, first with 5 mL pipettes, then with fire-polished glass Pasteur pipettes, and lastly with P1000 tips. Afterwards, cell suspension was filtered through a 70 *μ*M cell strainer before mixing with Debris Removal Solution. Cells were centrifuged after (300 g for 10 min) and washed with ice-cold FACS buffer (PBS, 2 % FBS, 0.78 mM EDTA). After centrifugation, cells were incubated for 30 min at 4 °C with FACS buffer containing Fc block (BD Biosciences) and primary antibody mix. Finally, right before loading the cells on the flow cytometer, cells were stained with DAPI (1:10000).

### Primary microglia isolation and culturing

For generation of microglial primary cultures, P0 WT or *Spp1*^*KO/KO*^ mice were decapitated, brain dissected from the skull and meninges removed in ice-cold HBSS with 5% FBS. 8 to 10 mice were pooled per culture preparation. Tissue was homogenized first with 2 mL pipettes (15 strokes) in 15 mL falcon tube and subsequently transferred to a pre-wet 50 mL tube with 70 *μ*M strainer. The 15 mL tube was washed with HBSS then put through filter to ensure all tissue was collected. The supernatant was removed and the cell pellet was resuspended in ice-cold 35 % isotonic percoll. The interface was carefully created with HBSS. The samples were centrifuged for 40 min at 4°C at 2800 g with no break and with slow acceleration and deceleration. The myelin layer and supernatant layers were aspirated, and the cell pellet was washed in HBSS. Cells were centrifuged and resuspended in 1 mL microglial media (DMEM F12 Gibco, 5 % fetal bovine serum (Gibco), 1 % pen-strep (Gibco), 50 ng/mL CSF1 416-ML-010/CF (RnD Systems), 50 ng/mL TGFb1 7666-MB-005/CF (RnD Systems), 100 ng/mL CX3CL1 472-FF-025/CF (RnD Systems) for cell counting. Cells were used within 7-10 days of plating.

### Isolation of synaptosomes and conjugation to pHrodo

3-5 8 week-old animals were pooled per experiment. In brief, mice were intra-cardiac perfused with 10 mL ice-cold PBS. The hippocampi and cortices were dissected on ice. Tissue was weighed and homogenized in 5 volumes of sucrose homogenization buffer (5 mM HEPES pH 7.4, 320 mM sucrose,1 mM EDTA) using a Dounce homogenizer with 15-20 strokes. The homogenate was centrifuged at 3,000 g for 10 min at 4°C and the supernatant was saved as total homogenate fraction (THF). The THF was centrifuged again at 14,000 g for 12 min at 4 °C and supernatant was saved as cytosolic fraction. The pellet was carefully resuspended in 550 *μ*l of Krebs-Ringer buffer (KRB: 10 mM HEPES, pH 7.4, 140 mM NaCl, 5 mM KCl, 5 mM glucose, 1 mM EDTA) and 450 µl of Percoll solution (for a final concentration of 45 %). The solution was mixed by gently inverting the tube and an interface was slowly created with 400 µl of KRB. After centrifugation at 14,000 g for 2 min at 4°C, the synaptosomal fraction was recovered at the surface of the flotation gradient and carefully re-suspended in 1 mL of KRB to wash. The functional synaptosomal preparation was centrifuged at 14,000 g for 1 min at 4 °C, after which the pellet was re-suspended in an appropriate volume of buffer. Fresh mouse synaptosomes were immediately divided into Eppendorfs at 2-2.5 mg of protein and re-suspended in total 1 mL KRB in 50 nM of oAβ 40-S26C dimer or just buffer and PBS as control and left overnight at 4°C on nutator. Synaptosomes were then centrifuged at 14,000 g for 1 min at 4°C, supernatant was discarded and synaptosomes were washed in 1 mL PBS, after which they were centrifuged at 14,000 g for 1 min at 4 °C to obtain oAβ-synaptosomes and control-synaptosomes. Briefly, 1 mg of synaptosomes were left at RT on nutator for 2 hr in sodium bicarbonate 0.1M with pHrodo(tm) Red, succinimidyl ester (P36600) at a concentration of 1 mg/mL. After conjugation, synaptosomes were centrifuged at 14000 g for 1 min, and resuspended for use in the *in vitro* synaptosome engulfment assay.

### *In vitro* synaptosome engulfment assay

Primary mouse microglia were treated with 1 µg of pHrodo-conjugated S26C oAβ-treated synaptosomes versus PBS control treated synaptosomes (PBS synaptosomes). For preferential engulfment studies, 1 *μ*g of both control and S26C oAβ-treated synaptosomes were added to the same well in the microglial media. Plates were then placed in a CD7 with the incubator at 37 °C and 5 % CO_2_. Fluorescent (594 nm and 647 nm) and brightfield (oblique and phase) images were acquired at a x20 objective (x0.5) at intervals of 3-5 min. A 3-slice z-stack was taken at 1.5 *μ*m interval to ensure that imaging was within focus throughout the imaging session however, one plane was used for analysis. For analysis, the z profile axis was plotted for respective pHrodos on ImageJ with respect to time. Fluorescence intensity at t=0 was subtracted from subsequent time frames.

### 3D-*τ*-STED/ STED-FLIM

For visualization of secreted SPP1 in mouse and human tissue, tissue sections were imaged with Leica STELLARIS 8 Stimulated Emission Depletion (STED) microscope using the 100x objective (1.4NA oil) (Leica Microsystems). Tissue sections were imaged at least 24 h after being coverslipped with mounting medium to avoid discrepancies in fluorescence lifetime within each section. STED microscope with Fluorescence Lifetime Imaging (STED-FLIM) was used to visualize secreted SPP1 in the extracellular space and fluorescence lifetime information was used to gate fluorescence signals. Alignment between STED laser and excitation laser was performed before each imaging session. 775 nm STED laser was used to generate a doughnut beam to silence the peripheral fluorophores from Alexa Flour 647 photoexcitation to achieve subdiffractional resolution. For all STED images, pixel size was limited to at most 50 nm. STED laser intensity was set at 20 %. All images were taken with a step size of 0.15 *μ*m. Fluorescence signal was time gated from -0.5 ms to 4.5 ms.

### Volume correlative light and electron microscopy (vCLEM) with Array Tomography

Mice were perfusion fixed with 4% PFA (EM grade) in ice-cold PBS as described above, the brain dissected and left in 4% PFA in PBS overnight at 4°C. The following day, coronal vibratome slices (100 *μ*m) of the brain were collected and slices containing clear cross sections of the hippocampus were manually trimmed to minimally contain the hippocampus and ensure that the tissue piece was asymmetric. Low resolution maps of the entire hippocampus tissue piece were taken using a confocal Zeiss LSM800 and 10x lens with montaging for gross mapping and identification of regions of interest. High resolution (63x 1.4-NA) confocal stacks were taken of regions of interest. Slices were fixed further with 2 % formaldehyde/ 1.5 % glutaraldehyde in 0.1M sodium cacodylate, before being processed for volume EM using a modified protocol based on the NCMIR protocol^76^ Briefly, tissue slices were incubated in 1 % Osmium tetroxide/ 1.5 % potassium ferricyanide for 1 h at 4°C, before being washed and left in 0.1M sodium cacodylate in a fridge overnight. The following day, tissue was incubated in 1% thiocarbohydrazide for 15 min at 60°C, 2 % osmium tetroxide for 30 min, 1 % uranyl acetate for 30 min and Walton’s lead aspartate for 30 min at 60°C, with numerous distilled water washes between each step. The samples were consequently dehydrated through an ethanol series, embedded in Epon resin and baked overnight at 60°C. Using the maps acquired by light microscopy, the block was trimmed down to the approximate region of interest and serial sections were collected on ITO coated coverslips using an ultramicrotome (Leica) and diamond knife (Diatome). Coverslips were mounted on SEM stubs using carbon stickies and silver DAG and array tomography serial backscattered electron images (5nm pixels) were acquired for each CLEM cell (n=4) using Atlas 5 software and a Gemini 300 SEM (Zeiss) operating in high vacuum, at 4.5kV with tandem decel operating at 3kV. Serial images were registered using TrakEM2^77^ in Fiji (NIH), aligned with the confocal images in Photoshop (Adobe) using nuclei as unbiased fiducials and 2/3D reconstructed using Amira (Thermofisher).

### scRNA-seq library preparation, expression and NicheNet analysis

Single cell libraries were prepared using 10X Genomics Chromium Next GEM Single Cell 3’ kit v3.1, according to the manufacturer’s instructions. As sorted PVMs and fibroblasts were in low abundance, we processed the whole sample, while for microglia cells we aimed for 5,000 cells per sample. Libraries were sequenced using an SP flowcell on an Illumina NovaSeq S6000 instrument, with sequencing parameters recommended by 10X Genomics, aiming for 40,000 reads per cell. Raw FASTQ files were pre-processed using 10X Genomics CellRanger (version 6.0) and the mm10 mouse reference genome. Filtered gene expression matrices obtained from CellRanger were imported in R (version 4.1.2) and analyzed using the Seurat (version 4.0.6) and NicheNet (version 1.0.0) packages^41^. Cells with more than 10% of reads aligning to mitochondrial transcripts were deemed to be low quality or damaged, and removed from the dataset. Identification of high variable genes, PCA and clustering were performed using Seurat functions. Cluster annotation was performed considering expression of a set of known cell type markers (Suppl Fig. 5). Differential expression analysis between pairs of conditions was performed with Seurat’s FindMarkers function using the Wilcoxon Rank Sum test. Comparisons between selected cell populations were performed using NicheNet functions and standard workflow.

### Statistics

All statistics analysis was performed in Prism (GraphPad Software, Version 9.3.1). Grubbs’ test (alpha=0.05) test was used to identify and discard outliers. Two groups were compared using two-tailed unpaired Student’s t-test. For oAβ injections experiments, two-tailed paired Student’s t-test were used to normalize variances of injections in different days or different oAβ batches. To compare more than 2 groups (WT, *Spp1*^*KO/KO*^, *App*^NL-F/NL-F^, *App*^NL-F^.*Spp1*^*KO/KO*^ mice or PVM, microglia and PVF), One way ANOVA with Kruskal-Wallis post hoc test was used. To compare multiple variables (WT and *Spp1*^*KO/KO*^ injected with S26C oAβ), two-way ANOVA with Tukey’s post hoc test was used. Data in graphs are presented as Mean ± S.E.M.

## Supplementary Information Titles and Legends

**Supplementary Figure 1. Related to Figure 1.**
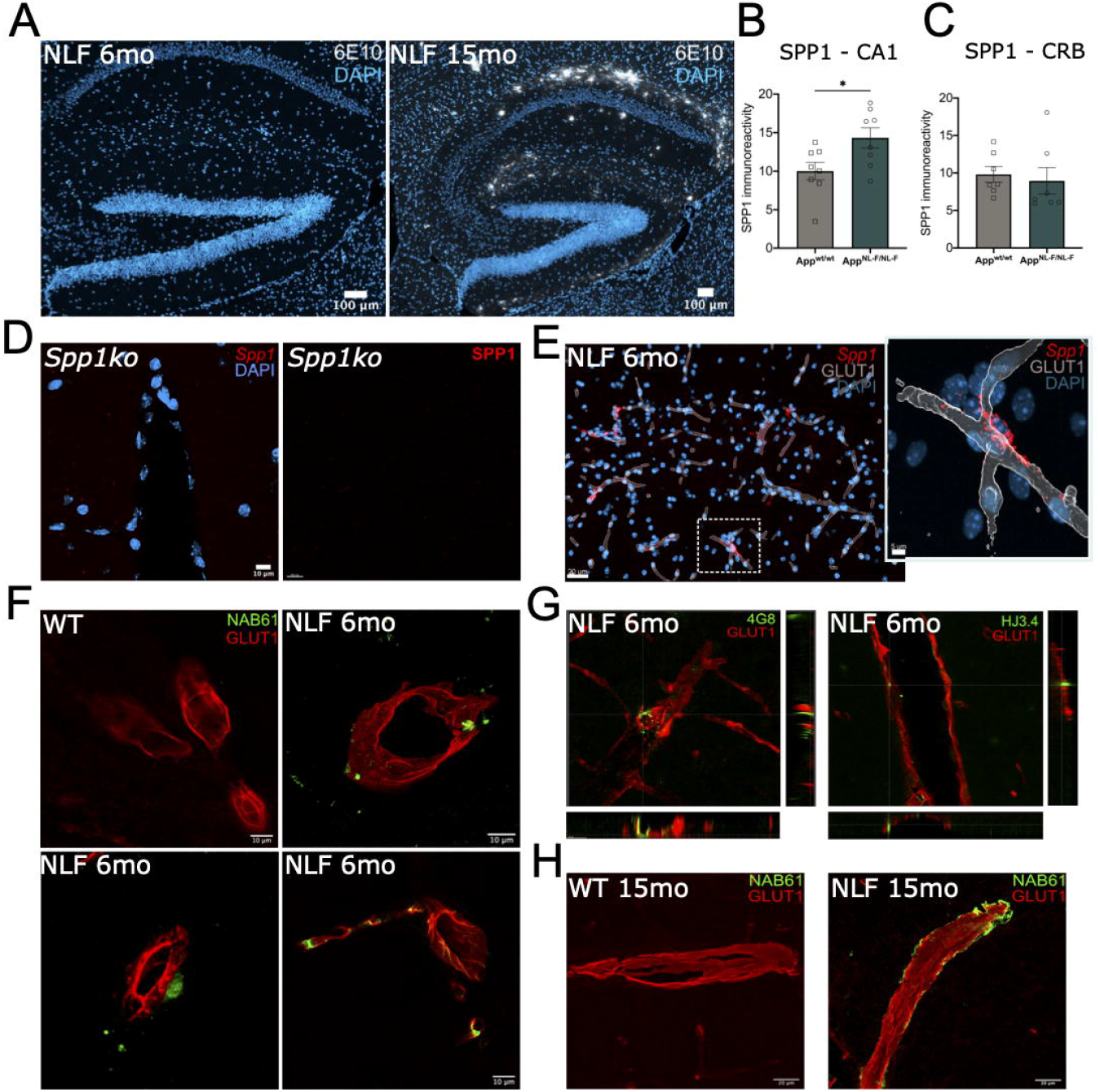
**(A)** Representative confocal images showing 6E10 plaque staining in hippocampus of 6 mo versus 15 mo *App*^NL-F^ mice. Scale bar represents 100 µm. **(B-C)** Quantification of SPP1 immunoreactivity in CA1 hippocampus **(B)** versus cerebellum as control region **(C)**, as measured by confocal imaging. Mann-Whitney test, n=8, 2 independent experiments. *p < 0.05. Data are shown as Mean ± SEM. **(D)** Validation of SPP1 antibody (left) and anti-*Spp1* probe for smFISH (right) in 3 mo *Spp1*^*KO/KO*^ hippocampal tissue. Scale bar represents 10 µm. **(E)** Representative confocal image of *Spp1* mRNA expression along GLUT1^+^ vasculature in SLM of 6 mo *App*^NL-F^ mice as characterized by smFISH-IHC. Scale bar represents 20 µm. Insert: 3D reconstruction. Scale bar represents 5 µm. Data are representative of at least 4 independent experiments. **(F)** Representative confocal images showing oAβ around GLUT1^+^ blood vessels as characterized by NAB61 immunoreactivity in 6 mo WT versus 6 mo *App*^NL-F^ SLM. Scale bar represents 10 µm. **(G)** Representative confocal images showing vascular oAβ deposition using alternative antibodies 4G8 and HJ3.4 that recognize Aβ17-24 and Aβ1-13, respectively, in 6 mo *App*^NL-F^ SLM. **(H)** Representative confocal images showing vascular oAβ (NAB61) in 15 mo WT versus *App*^NL-F^ SLM. Scale bar represents 20 µm.

**Supplementary Figure 2. Related to Figure 2.**
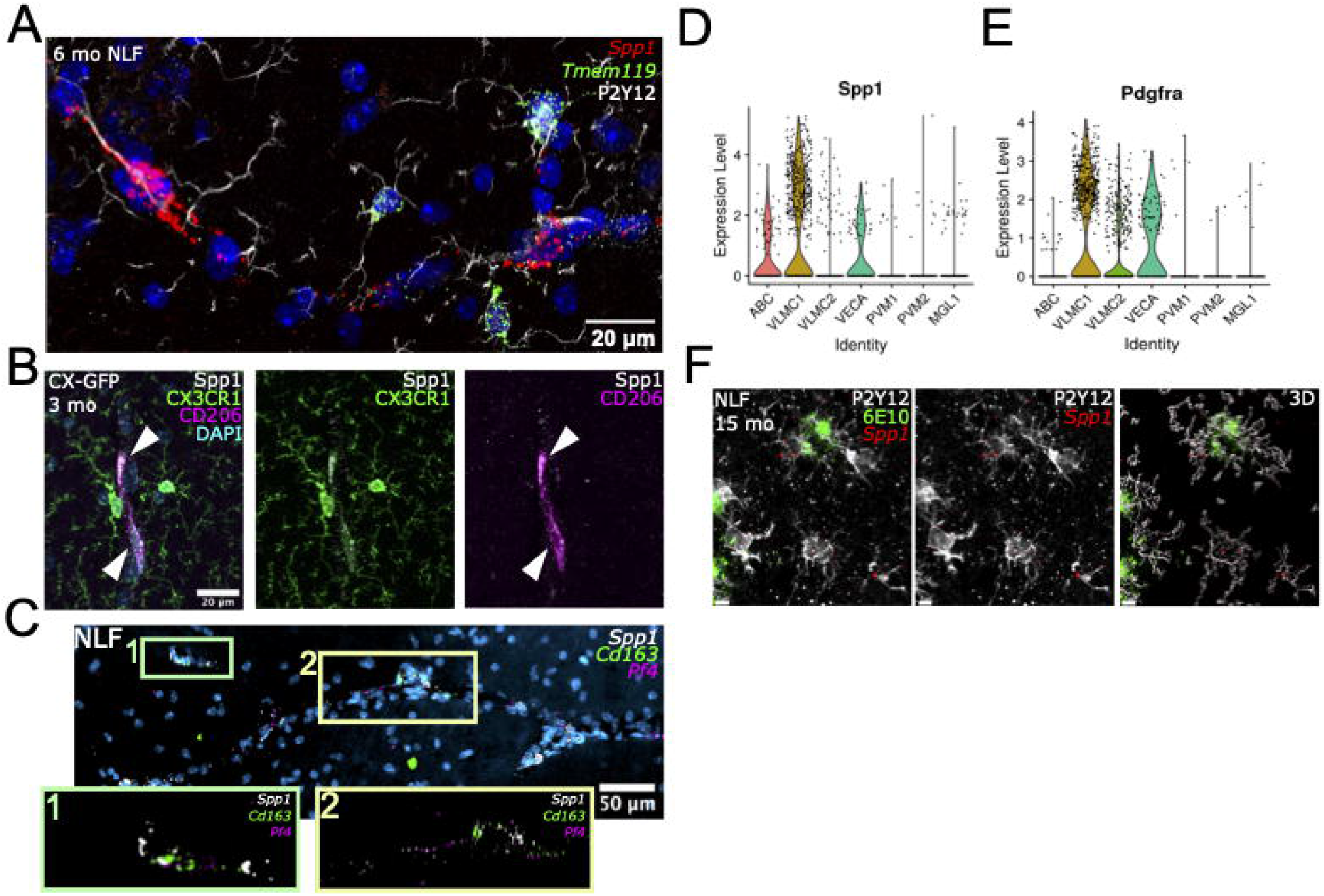
**(A)** Representative confocal image showing *Spp1* and *Tmem119* mRNA expression within P2Y12^+^ microglia of 6 mo *App*^NL-F^ SLM as characterized by smFISH-IHC. Scale bar represents 20 µm. **(B)** Representative confocal images of SPP1 protein expression in CD206^+^ PVM (arrow head) and CX3CR1^GFP^ myeloid cells of 3 mo *Cx3cr1*^*GFP/WT*^ SLM. Data are representative of at least 3 independent experiments. Scale bar represents 20 µm. **(C)** Representative confocal image showing *Spp1* mRNA expression occasionally colocalizing with pan-PVM markers *Cd163* and *Pf4 in* 6 mo *App*^NL-F^ SLM as identified by smFISH. Scale bar represents 50 µm **(D-E)** Violin plots of *Spp1* **(D)** and *Pdgfra* **(E)** expression reanalyzed from Zeisel et al., 2018. VLMC, vascular leptomeningeal cells; ABC, Arachnoid barrier cells; VECA, Arterial vascular endothelial cells; PVM, Perivascular macrophage; MGL, Microglia. **(F)** Representative confocal image showing *Spp1* mRNA in P2Y12 microglia associated with 6E10^+^ plaques in 15 mo *App*^NL-F^ SLM. Data are representative of 2 independent experiments. Scale bar represents 5 µm.

**Supplementary Figure 3. Related to Figure 2.**
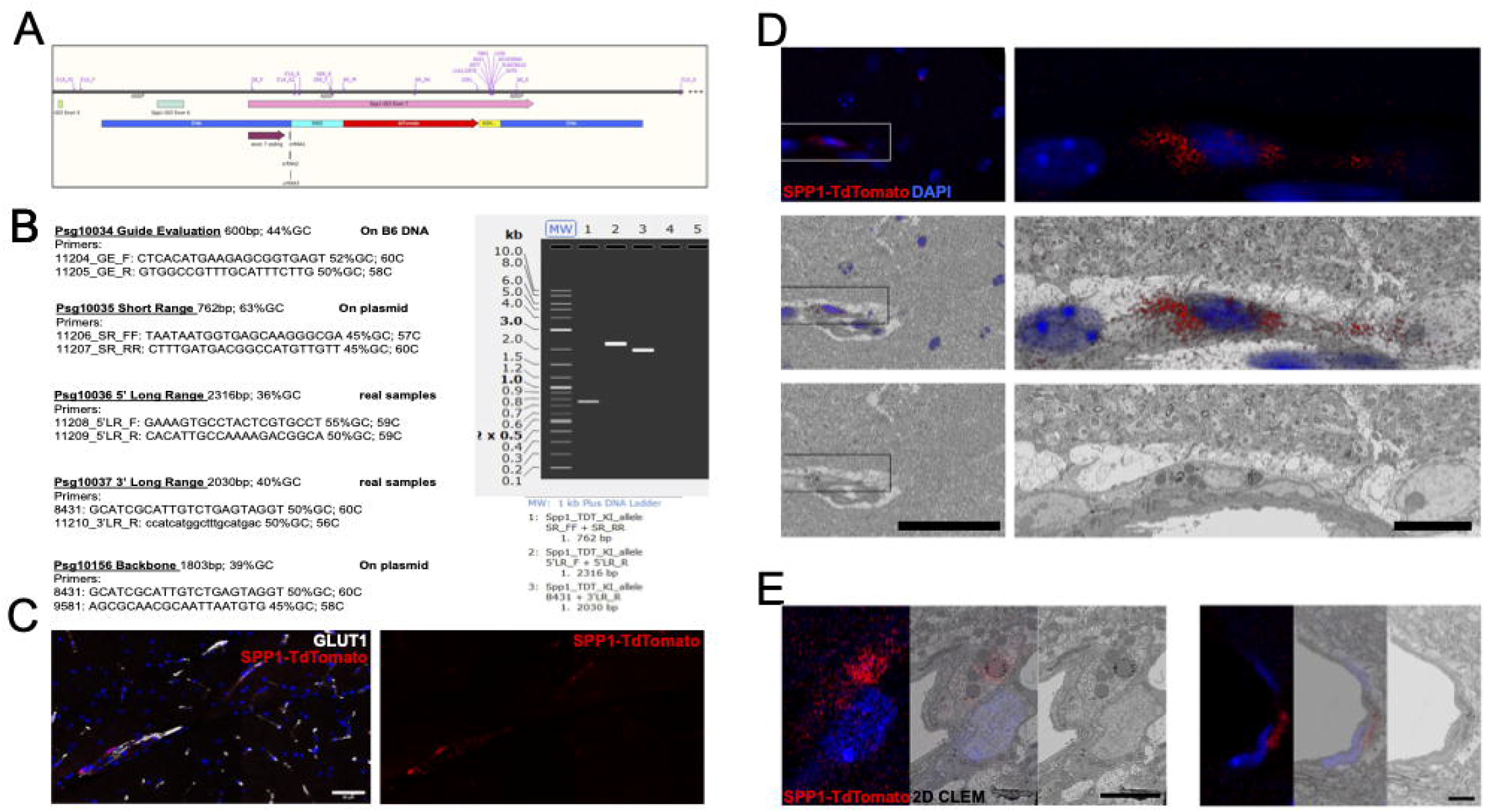
**(A)** Design of the knock-in allele is shown, with an IRES and TdTomato (TdT) reporter inserted at the stop codon in exon 7 in the mouse *Spp1* locus. Top, the mouse locus with locations of primers used for validation of the inserted construct and later genotyping for the reporter construct is shown. Below, schematic displays the upstream (5’HA) and downstream (3’HA) homology arms in blue; IRES in aqua; TdT reporter gene in red; and bovine growth hormone poly(A) region in yellow. **(B)** PCR primers used to confirm incorporation of construct are listed, with primer names based on genomic locations as shown in B. Gel shows appropriate size bands for incorporation of plasmid construct (lane 1), for left arm integration event (lane 2) and right arm integration event (lane 3). Primer pairs for plasmid backbone alone are negative (lanes 4 and 5). **(C)** Representative confocal image showing SPP1-TdT expression along GLUT1^+^ vessels in hippocampus of 3 mo *Spp1*^TdT^ reporter mice. **(D)** Left panels show low magnification overview of SPP1-TdT^+^ cell (red), and all nuclei (blue) in the hippocampus, as shown in Fig. 2H, scale bar 50 µm. Top, maximum intensity projection of confocal stack; middle, maximum intensity projection shown at reduced opacity over the correlated back scattered electron image of serial section 79 of the same region of the CLEM sample; bottom, back scattered electron image of serial section 79 of the same region of the CLEM sample alone. Box highlights region acquired across 200 serial sections. Right panels show boxed region at higher magnification with serial section 85. Scale bar represents 10 µm. **(E)** Two additional CLEM examples of SPP1-TdT^+^ cells targeted and correlated with volume electron microscopy. Left panels show maximum intensity projection of confocal stack; middle panels, maximum intensity projection shown at reduced opacity over the correlated back scattered electron image of a central serial section of the same region of the CLEM sample; right panels, back scattered electron image of the same region of the CLEM sample alone. Scale bar represent 5 µm.

**Supplementary Figure 4. Related to Figure 3.**
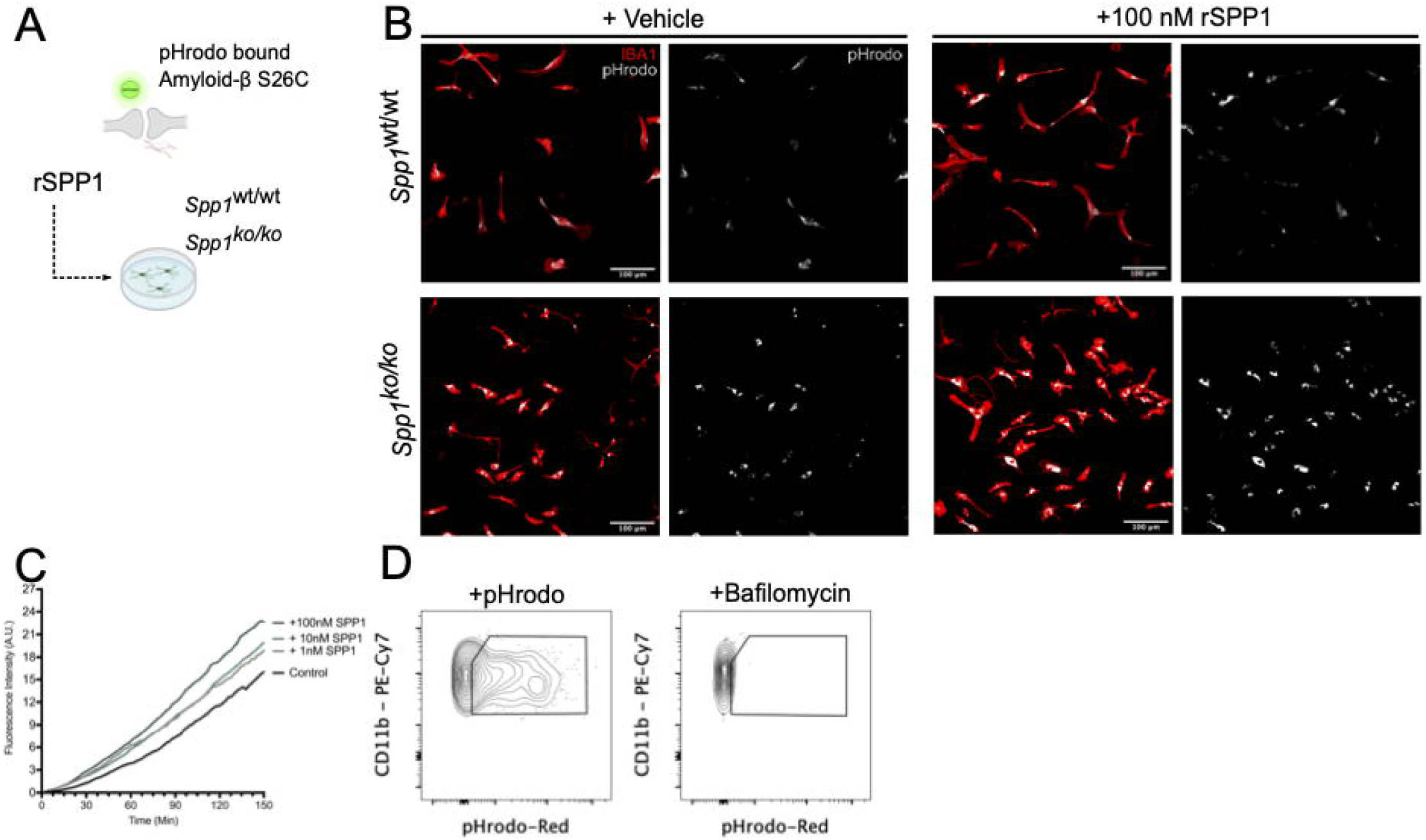
**(A)** Schematic illustrating *in vitro* engulfment assay whereby primary microglia isolated from WT or *Spp1*^*KO/KO*^ mice phagocytose S26C oAβ-treated synaptosomes that are tagged with pHrodo upon PBS or recombinant SPP1 treatment. **(B)** Representative images showing engulfment of pHrodo-synaptosomes by primary microglia isolated from WT mice (upper panel) or *Spp1*^*KO/KO*^ mice (lower panel), treated with vehicle (left) or 100 nM recombinant SPP1 (right). Data are representative of 2 independent experiments. Scale bar represents 100 µm. **(C)** Graph showing fluorescence intensity of pHrodo within primary WT microglial lysosomes over time (min), either in the presence of control or 1, 10 and 100 nM recombinant SPP1. **(D)** Representative FACS plot showing pHrodo signal in CD11b^+^ cells collected from primary microglial culture (left). No pHrodo signal was observed after Bafilomycin treatment (right). Data are representative of 2 independent experiments.

**Supplementary Figure 5. Related to Figure 5.**
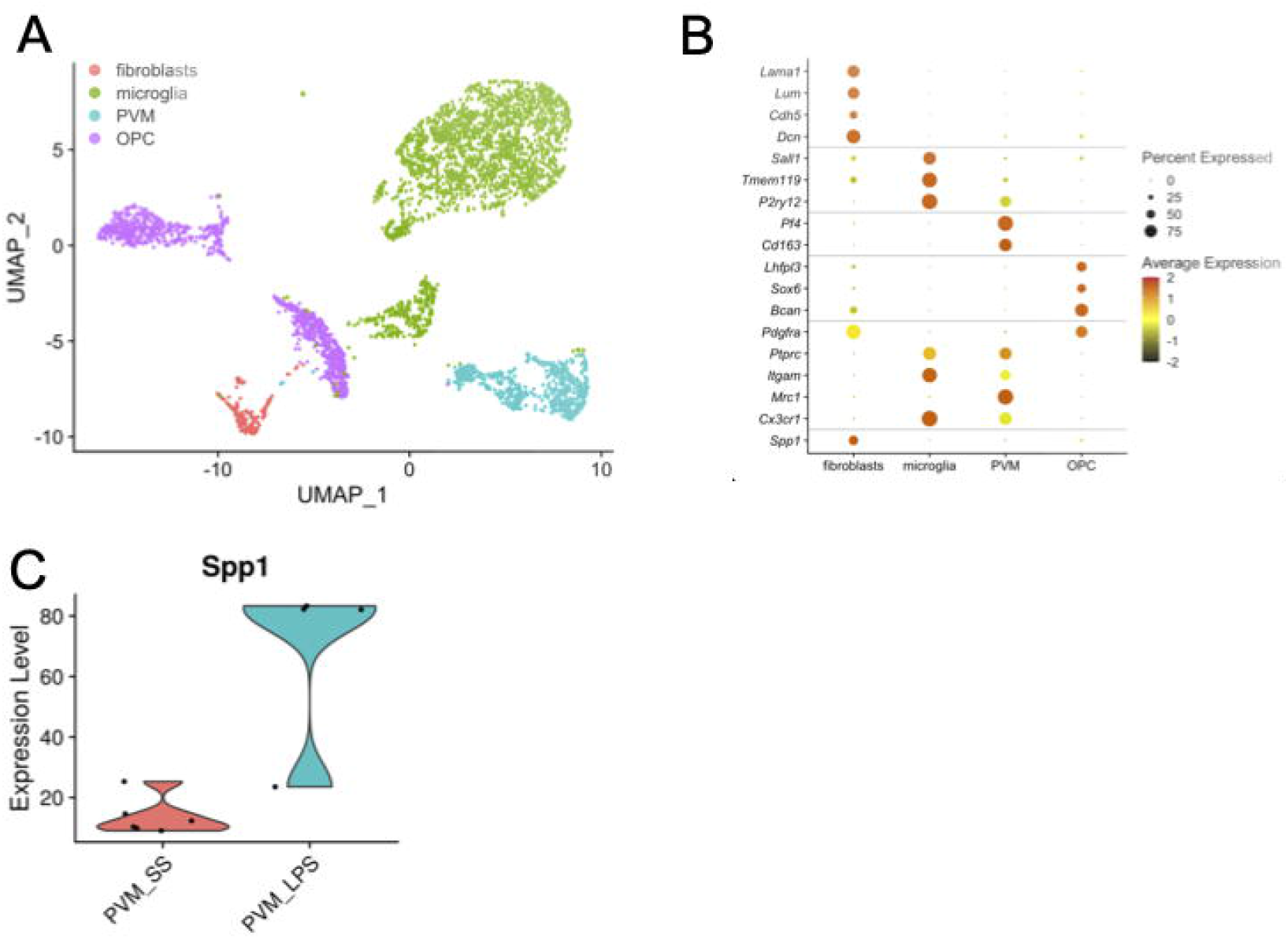
**(A)** UMAP plot of 4,238 cells, colour-coded based on the inferred cell type. **(B)** DotPlot of gene expression for a series of known cell type markers for fibroblasts (*Lama1, Lum, Cdh5, Dcn*), microglia (*Sall1, Tmem119, P2ry12*), PVM (*Pf4, Cd163*), and OPC (*Lhfpl3, Sox6, Bcan*). Expression of genes encoding for markers used for FACS of cells prior to scRNA-seq (*Pdgfra, Ptprc, Itgam, Mrc1, Cx3cr1*) and for *Spp1* is also included. Radius of dot is proportional to the percentage of cells expressing the gene, colour is the scaled gene expression level. **(C)** Violin plot of *Spp1* expression reanalyzed from Jung-Seok Kim et al., 2020^5^.

**Supplementary Table 1**

Case demographics for control and AD cases used in this study.

**Supplementary Movie 1**

Movie showing backscattered electron images of serial sections of SPP1-TdT PVM shown in Fig. 2H. The manually segmented nuclei of the SPP1-TdTomato positive cell is shown highlighted in blue, to mark the SPP1-TdTomato correlated cell. Movie width is 60 µm

